# The multi-stage plasticity in the aggression circuit underlying the winner effect

**DOI:** 10.1101/2024.08.19.608611

**Authors:** Rongzhen Yan, Dongyu Wei, Avni Varshneya, Lynn Shan, Hector J. Asencio, Dayu Lin

**Author notes:** These authors contributed equally to this work (Rongzhen Yan, Dongyu Wei).

## Abstract

Winning increases the readiness to attack and the probability of winning, a widespread phenomenon known as the “winner effect”. Here, we reveal a transition from target-specific to generalized aggression enhancement over 10 days of winning in male mice, which is supported by three stages of plasticity in the ventrolateral part of the ventromedial hypothalamus (VMHvl), a critical node for aggression. Over 10-day winning, VMHvl cells experience monotonic potentiation of long-range excitatory inputs, a transient local connectivity strengthening, and a delayed excitability increase. These plasticity events are causally linked. Optogenetically coactivating the posterior amygdala (PA) terminals and VMHvl cells potentiates the PA-VMHvl pathway and triggers the cascade of plasticity events as those during repeated winning. Optogenetically blocking PA-VMHvl synaptic potentiation eliminates all winning-induced plasticity. These results reveal the complex Hebbian synaptic and excitability plasticity in the aggression circuit during winning that ultimately leads to an increase in “aggressiveness” in repeated winners.

## Introduction

Aggression is an innate behavior across animal species. It is essential for competing for food, defending territory, securing mates, and protecting families and oneself. Across a wide range of species, aggression is expressed without learning. The neural circuit underlying aggression is believed to be genetically and developmentally hardwired (Lorenz, 1966; Tinbergen, 1951). However, the readiness to express aggression varies widely among individuals in the same species, even those with identical genetic backgrounds (e.g., inbred mice). The individual difference in aggression arises at least partly from prior experiences. Indeed, aggression is influenced by a wide variety of experiences, particularly winning and losing previous encounters (Hsu et al., 2006). In insects, reptiles, birds, and mammals, numerous studies have shown that winning experience leads to heightened aggression and an increased probability of winning, a phenomenon referred to as the “winner effect” (Hsu et al., 2006).

In the last decade, a few studies started to reveal the neural plasticity induced by winning and losing. Repeated winning in a tube test in mice enhanced the synaptic transmission from the medial thalamus to the medial prefrontal cortex, and suppressing this pathway blocked the winner effect in a tube test (Wang et al., 2011; Zhou et al., 2017). the synaptic connection from the medial thalamus to the medial prefrontal cortex was weakened in socially defeated animals (Franklin et al., 2017). In zebra fish, synaptic transmission in the lateral subregion of dorsal habenula decreased after defeat, and silencing this area eliminated the winner effect (Chou et al., 2016)

Recent studies further revealed synaptic plasticity in subcortical aggression-generating regions after winning. The ventrolateral part of the ventromedial hypothalamus (VMHvl) has now been firmly established as a critical region for aggression (Hashikawa et al., 2017; Lee et al., 2014; Lin et al., 2011; Yang et al., 2013). It is necessary, sufficient, and naturally active during aggressive behaviors in male and female mice. Nordman et al. found that transient potentiation of glutamatergic inputs from the medial amygdala posterior part (MeAp) to the VMHvl is responsible for aggression priming -- a short-term (within an hour) escalation of aggression after a brief exposure to a male conspecific (Nordman et al., 2020). Stagkourakis et al. found that the glutamatergic projection to the VMHvl from the posterior amygdala (PA), a region also functionally important for aggression, underwent synaptic potentiation after short-term winning (1-5 days)(Stagkourakis et al., 2020).

The studies above focused on the neural changes after one or a few victories. However, behavior studies revealed differential behavioral changes after short-term (∼ 3 days) vs. long-term winning (≥10 days). While both short- and long-term winning enhances aggression, as reflected by the increased attack probability and reduced attack latency, animals with 20 consecutive wins display aggressive behaviors even towards much heavier and stronger males or sometimes females (Kudryavtseva, 2012; Kudryavtseva et al., 2000; Kudryavtseva et al., 2014). The long-term winners also show less sensitivity to submissive signals from the opponent than short-term winners (Covington et al., 2019). Given the target- and context-unspecific attack, the long-term winners are sometimes considered “pathologically aggressive” (Kudriavtseva et al., 1997). Counter intuitively, some studies reported a decrease in the total attack time of long-term winners compared to short-term winners (Kudryavtseva et al., 2004). However, when the long-term winners (20 wins) are deprived of fighting opportunities, they show a “compensatory” increase in attack duration (Kudryavtseva et al., 2004). Given the distinct features of aggressive behaviors in short- and long-term winners, we hypothesize that the aggression circuits likely undergo different changes as the winning experience accumulates.

Our study aims to investigate changes in the aggression circuit over repeated winning. We first characterized aggressive behaviors in animals with short- and long-term winning experiences and then investigated synaptic and cellular changes in the VMHvl over the course of winning, with a particular focus on the PA input. Our results revealed multi-phased plasticity in the VMHvl over repeated winning, each with distinct temporal dynamics, that ultimately leads to heightened aggressiveness of an animal.

## Results

### Behavioral and physical changes induced repeated winning

We used repeated resident-intruder (RI) tests to provide winning experiences to the test mice. During the RI test, a non-aggressive group-housed Balb/C (BC) male mouse was introduced into the home cage of a single-housed C57BJ/6 male mouse for 10 minutes daily for up to 10 consecutive days **(Table S1)**. The BC intruder was randomly selected from a pool of 30 mice. If the resident male attacked the intruder and the intruder tried to escape and showed submissive postures, we considered the resident mouse aggressive and achieved a win. If the resident mouse closely investigated the intruder but did not attack during the 10-minute test, we considered the resident “non-aggressive” and underwent “social Interaction”. Only animals that consistently attacked and won across days (∼60% of all animals) were included in the final analysis. The 1-, 5- and 10-day winners refer to animals that have experienced 1, 5, and 10 consecutive wins, respectively (not including the current RI test if relevant) **(Figure 1A)**. “Naïve” mice are animals without RI test experience **(Figure 1A)**. Animals that showed no aggression during the 10-day RI tests constitute the “social” group.

**Figure 1.**
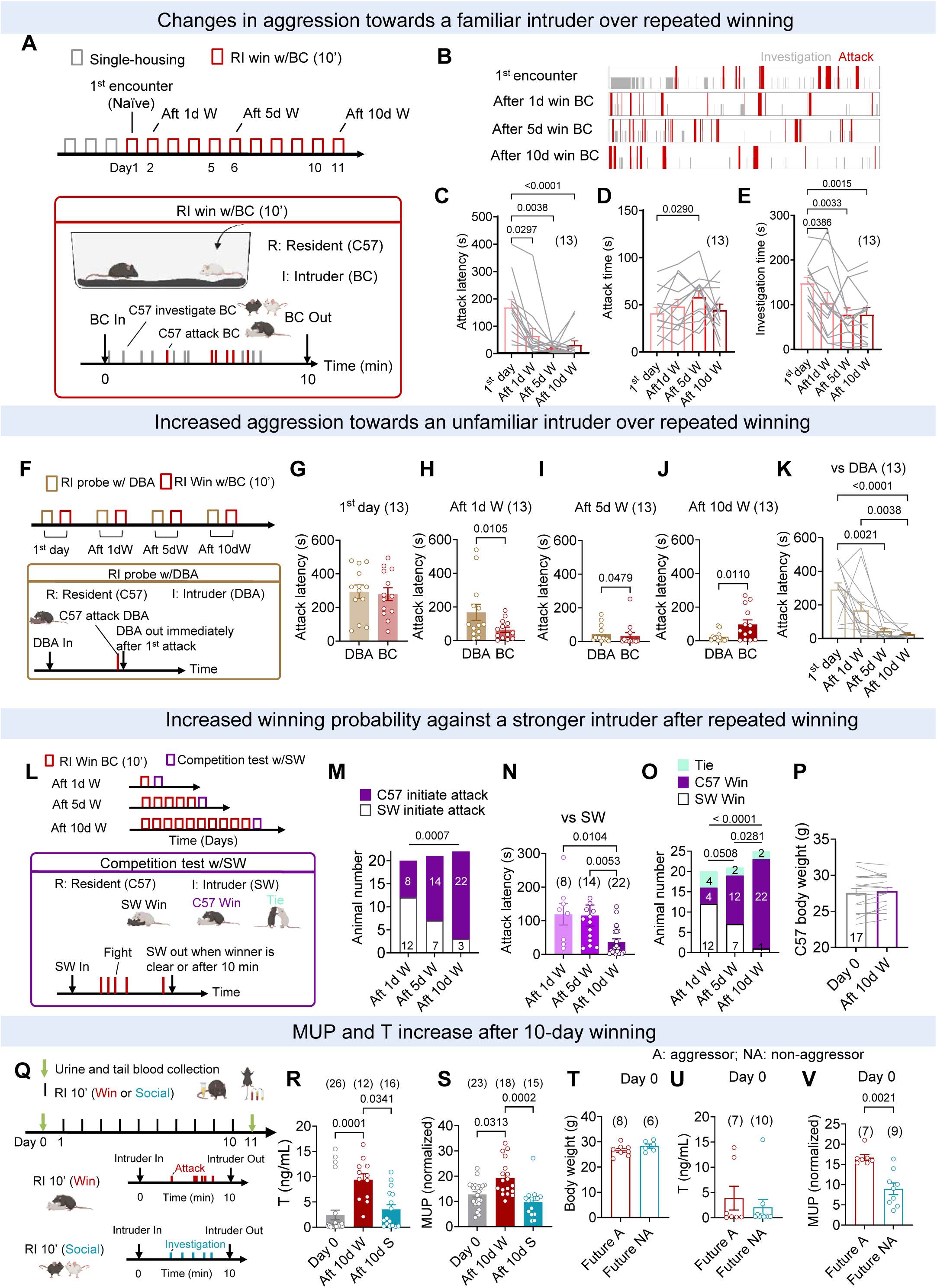
Changes in aggressive behaviors of male mice after repeated winning. (A) Illustration of the 10-day RI tests during which the test C57 residents repeatedly win the BC male intruders. (B) Raster plots showing attack and investigation during the RI tests. (C-E) Quantification of the latency to attack (C), attack duration (D), and social investigation duration (E) in mice with 0, 1, 5, and 10-day winning experiences. (F) Illustration of the probe test with DBA male intruders. (G-J) Latency to attack a BC and a DBA intruder in mice on the 1st day of RI test (G), after 1-day win with BC (H), after 5-day win with BC (I), and after 10-day win with BC (J). (K) Latency to attack DBA male intruders in mice with 0, 1, 5, and 10-day winning experiences with BC male intruders. (L) llustration of the competition test with SW male intruders. (M) Number of C57 and SW male mice that initiated the first attack in the competition test. (N) Latency to attack the SW intruder in mice with 1, 5, and 10-day wining experience with BC male intruders. (O) Number of C57 animals that win, lose and tie with the SW male intruder. (P) Body weight of C57 mice before and after 10 days of winning. (Q) Schematic representation of urine and blood collection in mice before and after 10-day win/social experiences with BC male intruders. (R-S) Quantification of testosterone (R) and MUP (S) levels in mice before and after 10-day win/social experiences with BC intruders. (T-V) Body weight (T), testosterone level (U), and MUP level (V) of naïve mice that attacked (future aggressors) and did not attack (future non-aggressors) during the RI test with BC male intruders on the day after the sample collection. Bars and error bars represent mean ± SEM. Circles and lines represent individual animals. Numbers in parenthesis or inside of bars indicate the number of subject animals. (C, K) Friedman test with repeated measures followed by Dunn’s multiple comparisons test. (D, E) One-way ANOVA with repeated measures followed by Tukey’s multiple comparisons test. (G, P) Paired t test. (H, I, J) Wilcoxon matched-pairs signed rank test. (M, O) Chi-square test. (R, S) Kruskal-Wallis test with Dunn’s multiple comparisons test. (T) Unpaired t test. (U, V) Mann-Whitney test. All statistical tests are two-tailed. All p:50.05 are specified. If not indicated, p> 0.05. See Supplementary Table 2 for detailed statistics. See also Figures S1 and S2 and table S1.

The aggression increased after 1-day winning, as indicated by the significantly shortened attack latency **(Figure 1B-C)**. After 5-day winning, the aggression against BC reached maximum, reflected by the lowest attack latency and the highest attack duration (**Figure 1C, D**). In 10-day winners, the aggression against BC appeared to decrease slightly from the 5-day winners, as indicated by a trend of increase in attack latency and decrease in attack duration, although overall, the aggression level remained high (**Figure 1C, D**). The decrease in aggression in 10-day winners may reflect increased familiarity with the BC intruder as the investigation duration gradually decreased over repeated RI tests **(Figure 1E)**.

The one-day winning-induced aggression increase is short-lived. While the attack latency decreased the next day after one day of winning, it returned to the initial level if tested a week later (**Figure S1A-E**). In contrast, the increase in aggression is highly stable in 10-day winners. After one week of single housing, the attack latency remained low, and the attack duration even increased, likely reflecting compensatory attack after aggression deprivation or decreased familiarity with the BC intruder **(Figure S1F-J)**(Kudryavtseva et al., 2014).

As the aggression level towards BC intruder over repeated winning could be affected by the simultaneous change in familiarity, we further probed the aggression level of naïve and 1-, 5- and 10-day winners using an unfamiliar non-aggressive DBA/2 male mouse (DBA) **(Figure 1F, Table S1)**. During the probe test, the DBA intruder was removed within 5 s after the resident initiated the first attack to minimize increases in familiarity **(Figure 1F)**. In naïve mice, we found that attack latency to BC and DBA was similar (**Figure 1G**). In 1-day winners (win over BC), the attack latency against BC mice was significantly shorter than that against DBA, suggesting the target-specific aggression increase in this early phase of winning **(Figure 1H)**. In 5-day winners (always win over BC), the latency to attack DBA and BC was similarly short **(Figure 1I)**. In 10-day winners (always win over BC), the attack latency towards the unfamiliar DBA was significantly shorter than that towards the familiar BC, supporting a suppression of aggression with increased familiarity (**Figure 1J**). When comparing DBA-directed aggression in animals with different days of winning experiences against BC, we found the attack latency to DBA was significantly shorter in 5- and 10-day, but not 1-day winners in comparison to naïve animals, supporting a transition from target-specific to generalized aggression increase over the course of repeated winning **(Figure 1K)**.

To further examine changes in “readiness to attack” and “winning probability”, two key parameters characterizing the winner effect, we challenged the C57 winners with aggressive, single-housed Swiss-Webster male intruders (SW) **(Table S1)**. The SW males are 40% heavier than the C57 intruders and will initiate attacks even as intruders, though often with a delay (Latency to initiate attack (Mean ± STD): 95.6 ± 33.7s). In this experiment, three different cohorts of C57 mice (1-, 5-, and 10-day winners against BC) were used. Each test mouse encountered the SW intruder only once and could win, lose, or tie with the SW **(Figure 1L)**. We found that as the C57 residents gained more winning experience, their readiness to attack strong SW intruders increased (**Figure 1M, N**). 10-day winners showed a significantly higher probability of initiating the 1^st^ attack after SW introduction and attacked with a shorter latency than other winner groups (**Figure 1M, N**). Importantly, repeated winning, even against a weaker intruder, increases the probability of winning against a stronger opponent (**Figure 1O**). 22/25 10-day winners defeated the SW intruder whereas only 4/20 1-day winners did so (**Figure 1O**). The animal’s body weight did not change significantly after the 10-day winning, suggesting that the increased winning probability was not due to increased physical advantage (**Figure 1P**).

Previous studies suggest repeated winning increases anxiety in CBA/Lac mice (Kudryavtseva et al., 2004). Interestingly, we observed an opposite behavior change in C57 10- day winners **(Figure S2A-I)**. In a light-dark box test, the distance and time spent in the light box gradually increased as the animals gained more winning experiences (**Figure S2B, F**). In contrast, repeated testing in single-housed or socially interacted mice minimally changed the performance in the light-dark box (**Figure S2C-D, G-H**). The increase in exploration in the light chamber is not due to a general increase in locomotion, as we did not observe significant experience-dependent changes in movement velocity when the animals were in their home cages or a large arena (**Figure S2J-O**). Thus, repeated winning, but not social interaction alone, increases adventurousness in C57 male mice, as reflected by their increased willingness to explore a risky environment.

Given the well-known role of circulating testosterone (T) in promoting male aggression (Barkley and Goldman, 1977; Dreher et al., 2016; Oyegbile and Marler, 2005) and the high levels of major urinary protein (MUP) found in dominant male mice (Lee et al., 2017), we measured the T and MUP levels in repeated winners (**Figure 1Q**). Indeed, both T and MUP concentrations were significantly higher in 10-day winners compared to naïve and 10-day social animals **(Figure 1R, S).** Some animals only engaged in social interaction and never became aggressive, while others became increasingly aggressive over repeated RI tests. We thus wondered whether any behavior or physiological parameters in naïve animals could predict future aggression. We found that future aggressors and non-aggressors did not differ in their body weight (**Figure 1T**), time spent, and distance traveled in the light box (**Figure S2E, I**), or testosterone level (**Figure 1U**). However, the MUP levels in the future aggressors and non-aggressors covered nearly non-overlapping ranges, with significantly higher values in future aggressors (**Figure 1V**). This data suggests that MUP level in naive mice could indicate an animal’s aggression potential while winning experience further increases MUP level and causes additional behavioral and physiological changes, especially aggression.

### Increase of VMHvl^Esr1^ cell responses to males over repeated winning

Changes in the neural circuits must support changes in aggressive behaviors. Given the critical role of estrogen receptor alpha-expressing VMHvl (VMHvl^Esr1^) cells in generating aggression (Hashikawa et al., 2017; Lee et al., 2014; Lin et al., 2011; Yang et al., 2013), we examined the responses of VMHvl^Esr1^ cells to aggression-provoking cues over repeated winning. We virally expressed GCaMP6f in VMHvl^Esr1^ cells using Esr1-2A-Cre male mice (Lee et al., 2014) and performed fiber photometry recording of the cell Ca^2+^ responses to anesthetized male and female BC mice and a toy mouse **(Figure 2A)**. To dissociate the behavioral and cell response changes due to winning, we presented the stimuli using a linear tracker when the recording mouse was head-fixed and awake (**Figure 2B**). Each animal was recorded before the first RI test and after 1-, 5- and 10-day winning or social interaction (**Figure 2C**).

**Figure 2.**
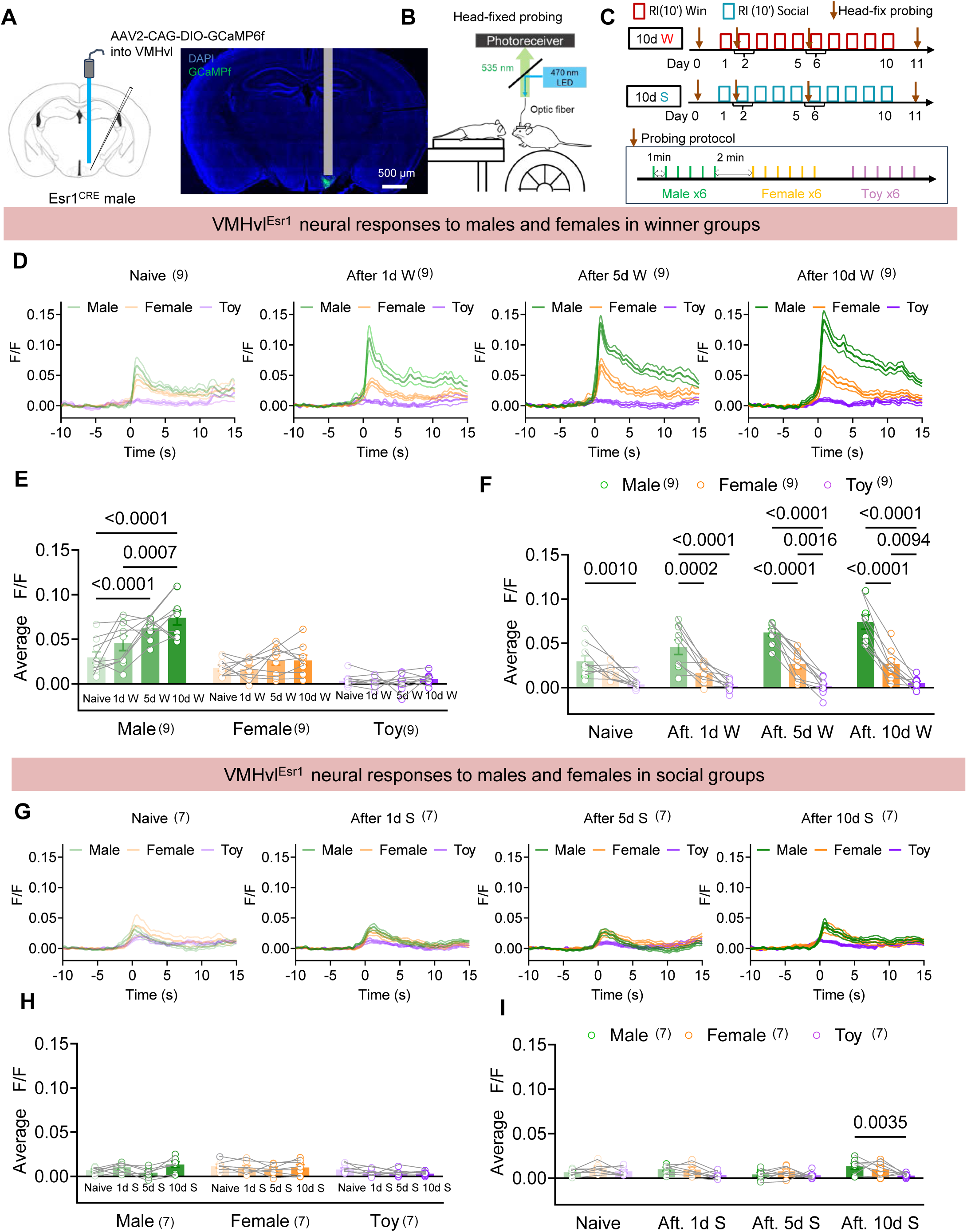
Change in VMHvl^Esr1^ cell responses to males over repeated winning and social interaction. (A) Virus injection, recording site and a representative histology image. (B) Schematic representation of the head-fixed fiber photometry recording setup. (C) Behavior paradigm and the probing protocol employed during head-fixed recordings. (D) Average peri-stimulus time histograms (PSTHs) aligned to the onset of male, female, and toy stimuli of all animals in the winner group after 0, 1, 5, and 10 days of winning. (E, F) Average GCaMP responses (LiF/F) during the presentation of male, female, and toy stimuli of all animals in the winner group after 0, 1, 5 and 10 days of winning. (G) Average PSTHs aligned to the onset of male, female, and toy stimuli of all animals in the social group after 0, 1, 5, and 10 days of social interaction. (H, I) Average GCaMP responses (LiF/F) during the presentation of male, female, and toy stimuli of all animals in the social group after 0, 1, 5, and 10 days of social interaction. Bar and error bar: mean ± SEM. Shades: ± SEM. Circles and lines represent individual animals. Numbers in parenthesis indicate the number of subject animals. (E, F, H, I) Two-way ANOVA with repeated measures followed by Tukey’s multiple comparisons test. All p :5 0.05 are specified. If not indicated, p> 0.05. See Supplementary Table 2 for detailed statistics.

The winning experience gradually increased the VMHvl^Esr1^ cell responses to BC males (**Figure 2D-F**). The VMHvl^Esr1^ cell response magnitude to BC male in 5- and 10-day winners was significantly higher than in naïve animals (**Figure 2D, E**). In contrast, VMHvl^Esr1^ cell response to females remained low after repeated winning (**Figure 2D, E**). Interestingly, naïve animals (before any RI test) did not show a difference in VMHvl^Esr1^ cell responses to BC males vs. females, although male mice nearly exclusively attack males (**Figure 2F**). After winning, the responses to males became significantly higher than those to females (**Figure 2F**). In contrast, the responses to both males and females remained low in non-aggressive animals after repeated social interactions (**Figure 2G, H**). Regardless of the winning or social interaction experience, VMHvl^Esr1^ cells showed minimum responses to the toy mouse, supporting the social-specific response patterns of the cells (**Figure 2D-I**).

### Changes in spontaneous synaptic transmission over repeated winning

The in vivo recording data supported the VMHvl as a site of change after winning. To understand whether the change occurs at the synaptic, cellular, or both levels, we performed in vitro patch clamp recording of VMHvl^Esr1^ cells using brain slices obtained from single-housed naïve, 1-, 5- and 10-day winners **(Figure 3A-C)**. We used Esr1-zsGreen transgenic male mice to facilitate the visualization of Esr1-positive cells (**Figure 3A**)(Saito et al., 2016). The recordings were performed the day after the last RI test **(Figure 3B)**.

**Figure 3.**
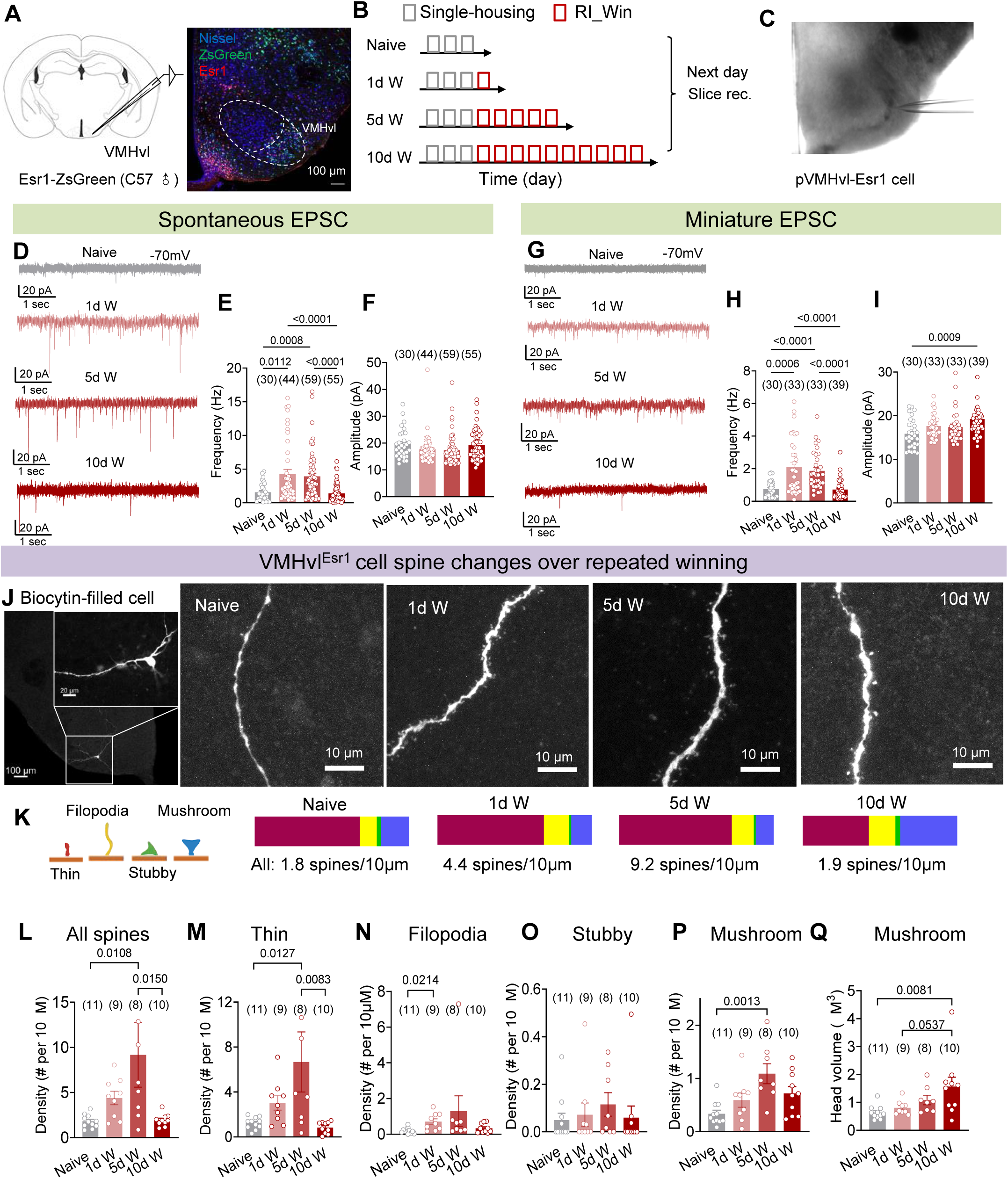
Synaptic transmission and spine morphology changes of VMHvl^Esr1^ cells over repeated winning. (A) Slice recording of VMHvl^Esr1^ cells and a representative histology image. (B) Experimental timeline. (C) A differential interference contrast (DIC) image showing a recorded cell in the VMHvl. (D) Representative voltage-clamp recording traces. (E, F) Quantification of the frequency (E) and amplitude (F) of sEPSCs of VMHVl^Esr1^ cells in mice with 0, 1, 5 and 10 days of winning experiences. (G) Representative voltage-clamp recording traces with 1µM TTX. (H, I) Quantification of the frequency (H) and amplitude (I) of mEPSCs of VMHVl^Esr1^ cells in mice with 0, 1, 5 and 10 days of winning experiences. (J) Representative image of a biocytin-filled VMHvl^Esr1^ cell (J1). Representative images of dendritic spine segmentations of mice with 0, 1, 5, and 10 days of winning (J2-J5). (K) Diagram depicting different types of spines (K1) and the proportion of each spine type in naïve mice and mice with 1, 5, and 10 days of winning experiences (K2-K5). (L) Total spine density in naïve mice and mice with different days of winning experiences. (M-P) The density of thin (M), filopodia (N), stubby (O), and mushroom (P) spines of VMHvl^Esr1^ cells of mice with 0, 1, 5, and 10 days of winning. (Q) Head volume of mushroom spines in mice with 0, 1, 5, and 10 days of winning experiences. Bar and error bar: mean ± SEM. Circles represent individual cells. Numbers in parenthesis indicate the number of cells. (E, F, H, I, N, O) Kruskal-Wallis test with Dunn’s multiple comparisons test. (L, M, P, Q) One-way ANOVA followed by Tukey’s multiple comparisons test. All p :5 0.05 are specified. If not indicated, p > 0.05. See Supplementary Table 2 for detailed statistics. See also Figs S3 and S4.

Consistent with a previous report (Stagkourakis et al., 2020), we observed a drastic increase in the frequency of spontaneous excitatory post-synaptic current (sEPSC) in 5-day winners compared to naïve animals (**Figure 3D, E**). This increase was also observed in 1-day winners, suggesting its rapid onset (**Figure 3D, E**). Surprisingly, the elevated sEPSC frequency could no longer be observed in 10-day winners (**Figure 3D, E**). In these animals, the sEPSC frequency was similar to that of naïve animals and significantly lower than 1- and 5-day winners (**Figure 3D, E**). To understand whether the change in sEPSC is spike-driven or not, we blocked action potentials using tetrodotoxin (1μM, TTX) and examined miniature EPSC (mEPSC) and found mEPSC frequency also increased in 1- and 5-day winners but not 10-day winners (**Figure 3G, H**). The amplitude of sEPSC did not change in any winning groups, whereas mEPSC magnitude significantly increased in 10-day winners (**Figure 3F, I**).

Additionally, we recorded spontaneous and miniature inhibitory post-synaptic currents (sIPSC and mIPSC) by holding the membrane potential at 0 mV. Unlike excitatory synaptic transmission, the frequency and amplitude in sIPSC did not differ between naïve and any winning groups, although the frequency of 10d win animals is lower than 5d win animals (**Figure SA-C**). The mIPSC frequency and amplitude did not differ across groups **(Figure S3D-F)**. These results suggest dynamic and complex changes in excitatory but not inhibitory synaptic transmission over repeated winning, featuring a transient increase in mEPSC frequency and a delayed increase in mEPSC amplitude.

### Spine morphology changes over repeated winning

Excitatory post-synaptic transmission occurs at dendritic spines, which are small protrusions from the dendrites (Yuste and Bonhoeffer, 2004). The density of VMHvl spines is known to be dynamically modulated (Calizo and Flanagan-Cato, 2000; Dias et al., 2021; Frankfurt et al., 1990; Stagkourakis et al., 2020). Thus, we wondered whether the up and down of m/sEPSC frequency over the course of winning can be explained by changes in spine density. We filled the recorded VMHvl^Esr1^ cells with biocytin and observed a drastic increase in the spine density in 1d and 5d winners, but not 10d winners, in comparison to naïve mice (**Figure 3J, K, L**), matching the time course of m/sEPSC frequency changes (**Figure 3E, H**).

We further characterized the spine morphology. Spines can be classified as filopodia, stubby, thin, and mushroom based on their shape and size (**Figure 3K**)(Ghani et al., 2017; Pchitskaya and Bezprozvanny, 2020). Filopodia spines are the most dynamic and may last only for minutes (Ziv and Smith, 1996). Thin and stubby spines could last several days (Holtmaat et al., 2005). Mushroom spines could be stable for months and mediate the largest synaptic current among all types of spines (Grutzendler et al., 2002). In naïve animals, the overall spine density was low (1.8 spines/10µm), and thin spines constituted 68% of all spines, followed by mushroom spines (18%), filopodia (11%) and stubby spines (3%) (**Figure 3K**). In 1- and 5-day winners, there was a gradual increase in the density of all types of spines (**Figure 3L-P**). But relatively speaking, thin spines and filopodia showed larger increases than others. In 5-day winners, thin spines and filopodia increased to 87% of all spines, while the fraction of mushroom spines decreased to 12% despite its net increase in number (**Figure 3K, P**). In 10-day winners, all spines showed a decrease in density, but mushroom spine density decreased the least (**Figure 4M-P**). In 10-day winners, thin spines and filopodia constituted approximately 60% of the total spine, while mushroom spines comprised 37%, doubling their share as in naïve animals (**Figure 4K**). Furthermore, we found that the head volume of mushroom spines increased gradually over repeated winning and almost tripled in size in 10-day winners compared to naïve animals (**Figure 4Q**). In contrast, we observed no change in spine density or morphology after 10-day social interaction compared to naïve animals, suggesting that winning experience is required to increase mushroom spine density and head volume (**Figure S4**). These results indicate that the increased m/sEPSC frequency after short-term winning is likely due to many newly emerged spines. Most of these new spines are unstable and disappear in 10-day winners, causing a decrease in m/sEPSC frequency, while the more stable mushroom spines survive and grow in size, likely contributing to the larger mEPSC amplitude in 10-day winners.

**Figure 4.**
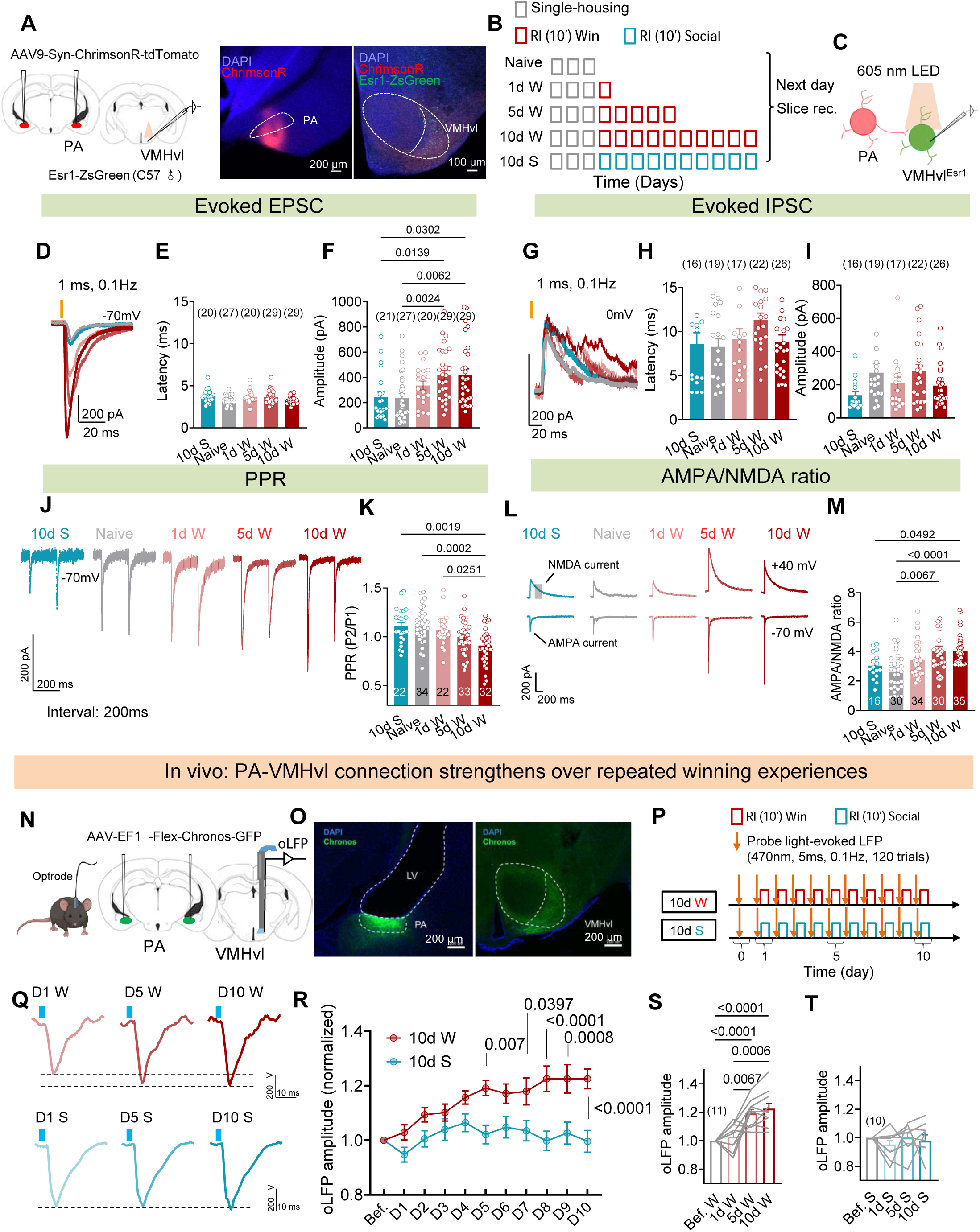
Monotonic potentiation of PA-VMHvl pathway over repeated winning. (A) Schematic of the virus injection, recording site and a representative histology image showing the expression of ChrimsonR and ZsGreen. (B) Experimental timeline. (C) Schematics of the slice recording of VMHvl^Esr1^ cell responses to PA terminal activation. (D) Representative traces of oEPSC of VMHvl^Esr1^ cells from male mice with 0, 1, 5, 10 days of winning experience and 10 days of social interaction. (E, F) Latency (E) and amplitude (F) of oEPSC of VMHvl^Esr1^ cells from male mice with 0, 1, 5, 10 days of winning experience and 10 days of social interaction. (G) Representative trace of oIPSC of VMHvl^Esr1^ cells from male mice with 0, 1, 5, 10 days of winning experience and 10 days of social interaction. (H, I) Latency (H) and amplitude (I) of oIPSC of VMHvl^Esr1^ cells from male mice with 0, 1, 5, 10 days of winning experience and 10 days of social interaction. (J) Representative oEPSCs after a pair of light pulses with 200 ms interval. (K) PPR of VMHvl^Esr1^ cells from male mice with 0, 1, 5, 10 days of winning experience and 10 days of social interaction. (L) Representative light-evoked AMPA and NMDA currents. (M) AMPA/NMDA ratio of VMHvl^Esr1^ cells from male mice with 0, 1, 5, 10 days of winning experience and 10 days of social interaction. (N) Diagram showing the in vivo optrode recording strategy. (O) Histological image showing the Chronos expression in the PA and VMHvl. (P) Timeline of behavior training and oLFP probing. (Q) Representative oLFP traces in animals undergone repeated winning or social interaction. (R) Amplitude of oLFP across days. The value is normalized to the before RI test level,. (S) Normalized oLFP amplitude over repeated winning experiences. (T) Normalized oLFP amplitude over repeated social interactions. Bar and error bar: mean ± SEM. Circles in E, F, H, I, K, M represent individual cells. Lines in S and T represent individual animals. Numbers in parenthesis or inside of the bar in E, F, H, I, K, M indicate the number of recorded cells. Numbers in parenthesis in S and T indicate the number of subject animals. (E, K) One-way ANOVA followed by Tukey’s multiple comparisons test. (F, H, I, M) Kruskal-Wallis test with Dunn’s multiple comparisons test. (R) Two-way ANOVA with repeated measures followed by Sidak’s multiple comparisons test. (S) One sample t test and one-way ANOVA with repeated measures followed by Tukey’s multiple comparisons test. (T) One sample Wilcoxon test followed by Holm-Sidak correction and Friedman test with repeated measures followed by Dunn’s multiple comparisons test. All statistical tests are two-tailed. All p :5 0.05 are specified. If not indicated, p > 0.05. See Supplementary Table 2 for detailed statistics.

### Increase in PA-VMHvl synaptic transmission over repeated winning

It is intriguing that some spines only transiently appear while others hang around for much longer over the course of winning. We next aimed to elucidate the presynaptic partners of these transient vs. stable spines. PA is a primary source of extra-hypothalamic excitatory input to the VMHvl, and a previous study found that the PA-VMHvl connection strengthened after 5-day winning (Stagkourakis et al., 2020). Thus, we asked whether the synaptic potentiation of the PA-VMHvl pathway disappears or maintains after 10-day winning. The former scenario will suggest PA as a source of input to the transient spines, while the latter will suggest PA may partner with the longer-lasting mushroom spines.

We first examined the PA-VMHvl synaptic connection *in vitro*. We virally expressed ChrimsonR in the PA glutamatergic cells of Esr1-zsGreen male mice and subjected the animals to single-housing alone, 1d, 5d, or 10d winning and 10d social interaction **(Figure 4A-B).** Four weeks after virus injection, we prepared the brain slices containing the VMHvl, performed the voltage-clamp recording of the zsGreen positive VMHvl cells, and examined the VMHvl^Esr1^ EPSCs evoked by 1ms light activation of PA terminals (**Figure 4C**). The latency of optogenetically evoked EPSC (oEPSC) was short in all groups (∼3.5 ms), suggesting the monosynaptic nature of the connection, as previously shown (**Figure 4D, E**) (Stagkourakis et al., 2020; Yamaguchi et al., 2020). However, the amplitude of oEPSC of VMHvl^Esr1^ cells varied significantly with animals’ experiences (**Figure 4F**). Compared to naïve animals, the oEPSC amplitude showed a trend of increase in 1-day winners and became significantly higher in both 5d and 10d winners (**Figure 4F**). In 10-day social animals, we observed no increase in oEPSC, suggesting the PA-VMHvl potentiation is winning-dependent (**Figure 4F**). In contrast to the oEPSC, light-evoked inhibitory post-synaptic current (oIPSC) amplitude did not change after winning **(Figure 4G, I)**. Consistent with our previous findings (Yamaguchi et al., 2020), the oIPSC has a long latency (∼9.3 ms) and thus is likely polysynaptic **(Figure 4G, H)**.

To understand whether the change in oEPSC occurs at pre- or post-synaptic sites, we measured the paired-pulse ratio (PPR), a parameter reflecting the presynaptic vesicle-releasing probability (**Figure 4J**). We found a gradual decrease in PPR in 1-, 5- and 10-day winners, indicating increased presynaptic release probability of PA cells over repeated winning (**Figure 4J, K**). In contrast, PPR in social animals did not differ from that in naïve animals **(Figure 4J, K)**. We further investigated potential changes in post-synaptic responses by measuring the relative synaptic currents mediated by α-amino-3-hydroxy-5-methylisoxazole-4-propionate (AMPA) and N-methyl-D-aspartate (NMDA) receptors. AMPA/NMDA ratio could reflect the composition of spine types as AMPA receptors are abundant in mushroom spines (up to 150/spine) but only sparsely present in thin and filopodia spines (Matsuzaki et al., 2001). In contrast, the NMDA receptor number does not differ drastically across spines (Matsuzaki et al., 2001). We measured the AMPA receptor-mediated current by holding the membrane potential at -70mV, a voltage preventing NMDA receptor from being activated due to Mg^2+^ blockage. The NMDA receptor-mediated current was measured at +40 mV (to unblock Mg^2+^) and 60-65 ms after the light onset, as AMPA receptor-mediated current decays quickly (Stagkourakis et al., 2020). The AMPA/NMDA ratio gradually increased with winning experience and is significantly higher in 5d and 10 day winners than in naïve animals (**Figure 4L, M**). In contrast, AMPA/NMDA ratio did not change after 10 days of social interaction **(Figure 4L, M).** These results suggest that PA-VMHvl synaptic potentiation likely involves both pre- and post-synaptic changes.

We further monitored the strength of the PA-VMHvl pathway in vivo over repeated winning by expressing Chronos in PA^Esr1^ cells and implanting an optrode in the VMHvl of Esr1-2A-Cre male mice (**Figure 4N, O**) (Klapoetke et al., 2014). Four weeks after surgery, we probed PA to VMHvl terminal stimulation-evoked local field potential (oLFP) by delivering 5 ms, 473 nm light pulses (0.1Hz, 120 times) to the VMHvl in naïve animals. We then probed oLFPs daily for 10 consecutive days (before the RI test) as the animals gained a daily winning experience (**Figure 4P**). We found that the oLFP amplitude increased gradually in winning mice over days but did not change in social mice that never attacked (**Figure 4Q-T**). These in vivo and in vitro recording results suggest that the synaptic transmission from PA to VMHvl cells is potentiated monotonically over repeated winning. Thus, PA inputs are likely to partner with the stable mushroom spines that emerge and grow over 10 days of winning experience.

### Transiently appeared intra-VMHvl synapses

Given that the VMHvl synapses with long-range PA inputs strengthen continuously over 10-day winning, we wonder whether the transiently appeared spines represent a qualitatively different type of connection, e.g., local connections among VMHvl cells. VMHvl^Esr1^ cells are overwhelmingly glutamatergic (Hashikawa et al., 2017) and have been suggested to form extensive local connections (Nishizuka and Pfaff, 1989). However, recent pair-recordings revealed surprisingly sparse intra-VMH connections in naïve animals (Shao et al., 2022).

To test this hypothesis, we expressed Cre-dependent hM4Di in the VMHvl^Esr1^ cells, subjected the animals to 0, 1, 5, or 10 days of winning, and performed patch clamp recording of VMHvl^Esr1^ cells on the day after the last win, which was approximately 3 weeks after virus injection (**Figure 5A, B**). During the recording, we added CNO, which is known to block the synaptic release of hM4Di-expressing cells (**Figure 5C**)(Stachniak et al., 2014), into the bath solution. Before the CNO application, as expected, we observed higher sEPSC frequency in the 1d and 5d winners than in naïve and 10d winners (**Figure 5D-I**). After CNO perfusion, the sEPSC frequency of 1d and 5d winners, but not naïve and 10d winners, significantly decreased (**Figure 5D-H**). After CNO-mediated suppression, we found no difference in sEPSC frequency across groups (**Figure 5K**). The sEPSC amplitude did not differ among groups regardless of CNO treatment (**Figure 5J, L**). These results suggest that first, the intra-VMHvl cell connection is likely minimal in naïve and 10-day winners as blocking synaptic inputs from neighboring VMHvl cells has no effect on sEPSC frequency, and second, the increased sEPSC frequency in 1d and 5d winners can be accounted largely by the enhanced synaptic transmission among VMHvl cells as blocking VMHvl inputs is sufficient to eliminate the increase.

**Figure 5.**
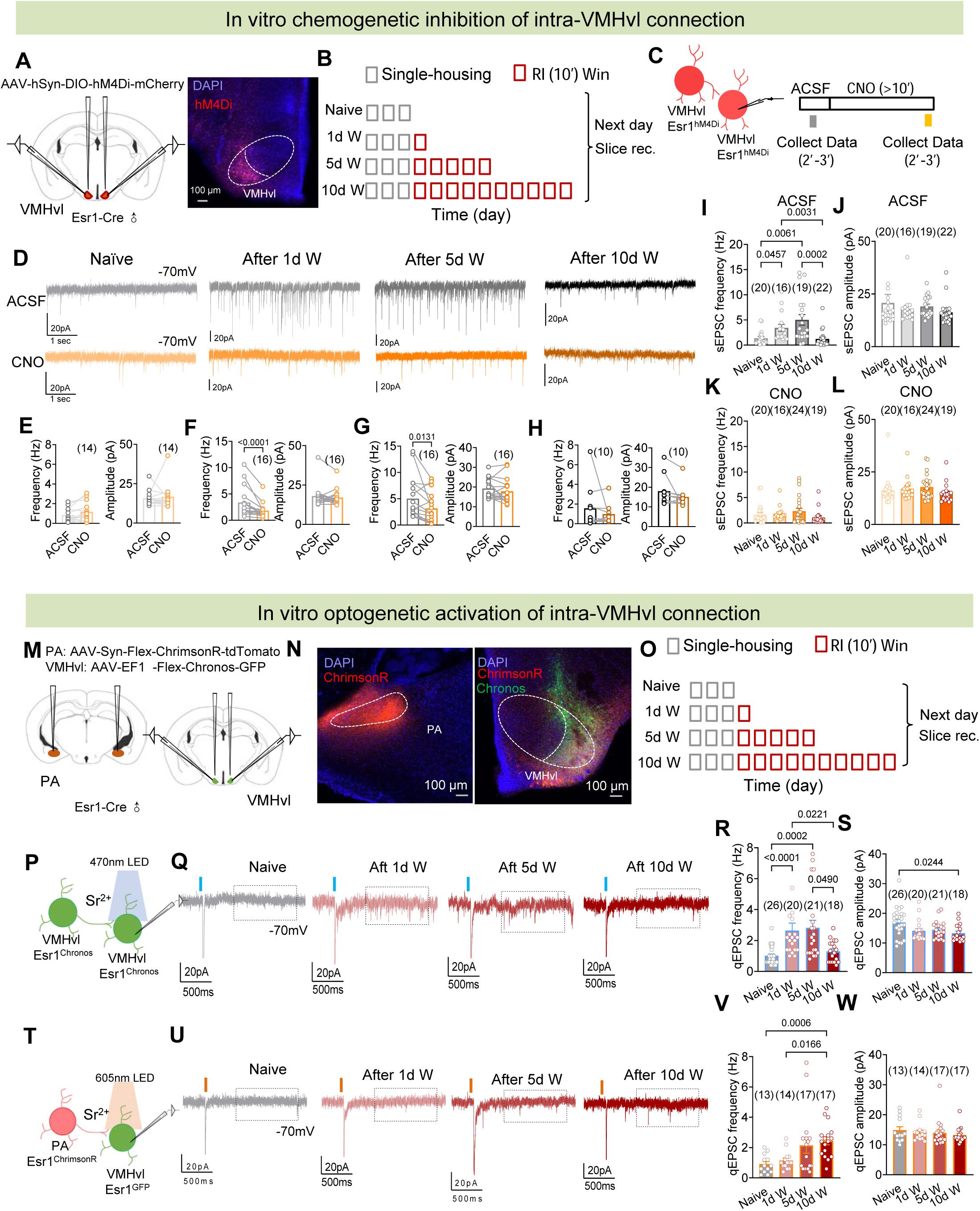
Transient strengthening of intra-VMHvl connectivity over repeated winning. (A) Virus injection, recording location and a representative histology image. (B) Experimental timeline. (C) Slice recording strategy to inhibit and probe intra-VMHvl connection. (D) Representative voltage-clamp recording traces before and after 10 µM CNO perfusion. (E-H) Quantification of the frequency and amplitude of sEPSCs before and after CNO application in naïve mice (E), 1-day winners (F), 5-day winners (G), and 10-day winners (H). (I-J) The frequency (I) and amplitude (J) of sEPSCs before CNO perfusion. (K-L) The frequency (K) and amplitude (L) of sEPSCs after CNO perfusion. (M) Virus injection, recording location and representative histology images. (N) Histological depiction of virus expression in PA and VMHvl. (O) Experimental timeline. (P) Slice recording strategy to optogenetically activate and record intra-VMHvl connection. (Q) Representative voltage-clamp recording traces after 1ms 470nm light activation of VMHvl^Esr1^ cells. (R-S) The frequency (R) and amplitude (S) of qEPSCs in response to VMHvl^Esr1^ activation in naïve mice and mice with 1-day, 5-day, and 10-day winning experiences. (T) Slice recording strategy to optogenetically activate PA terminals and record qEPSCs of VMHvl^Esr1^ cells. (U) Representative voltage-clamp recording traces after 1ms 605nm light activation of PA terminals. (V-W) The frequency (V) and amplitude (W) of qEPSCs in response to PA terminal activation in naïve mice and mice with 1-day, 5-day, and 10-day winning experiences. Bar and error bar: mean ± SEM. Circles and lines represent individual recorded cells. Numbers in parenthesis indicate the number of cells. (E, F, G, H) Wilcoxon matched-pairs signed rank test. (I, J, K, L, R, S, V, W) Kruskal-Wallis test with Dunn’s multiple comparisons test. All p :5 0.05 are specified. If not indicated, p > 0.05. See Supplementary Table 2 for detailed statistics.

Additionally, we optogenetically probed the intra-VMHvl connection in vitro in naïve male mice, 1d, 5d, and 10d winners (**Figure 5M-O**). Specifically, we expressed Chronos in VMHvl^Esr1^ cells and performed patch clamp recording of VMHvl^Esr1^ cells while optogenetically inducing glutamate release from presynaptic VMHvl^Esr1^ cells (**Figure 5P**). Importantly, we replaced Ca^2+^ with Sr^2+^ (4 mM) in the extracellular solution. Due to the low efficiency of Sr^2+^ in triggering synaptic vesicle release, we can isolate the currents caused by presynaptic inputs from direct Chronos activation by examining EPSCs occurred between 0.5-1.5s after light delivery (**Figure 5P, Q**) (Bekkers and Clements, 1999; Keppeler et al., 2018; Lee et al., 2023). As the light-evoked EPSC is asynchronous, each event is likely triggered by releasing a single presynaptic vesicle, namely quantal EPSC (Abdul-Ghani et al., 1996). To prevent the polysynaptic EPSCs, tetrodotoxin (1μM, TTX) and 4-aminopyridine (100 μM, 4AP) were added to the extracellular solutions (Petreanu et al., 2009). In naïve animals, we found the rate of light-evoked qEPSC was low (1.02 ± 0.11 event/light pulse), further supporting extremely sparse connection among VMHvl cells (**Figure 5Q, R**). In 1d and 5d winners, but not 10d winners, there was a significant increase in the qESPC frequency compared to naïve animals, supporting a transient increase in local VMHvl cell connectivity after short-term winning (**Figure 5Q, R**). We also noted that the amplitude of qEPSC in winner groups tended to be lower than in naïve animals, suggesting a lower glutamate load in the presynaptic ventricles of these newly formed synapses (**Figure 5S**).

For comparison, we injected Cre-dependent ChrimsonR into the PA in the same animals to record the VMHvl synaptic responses to quantal PA inputs (**Figure 6T**). In contrast to the VMHvl responses to the VMHvl inputs, we found the qEPSC frequency evoked by PA input gradually increased with winning experience, further supporting a monotonic increase in PA-VMHvl connection strength over the course of 10-day winning (**Figure 5U, V**). The amplitude to PA-evoked qEPSC of VMHvl^Esr1^ cells remained similar after winning (**Figure 5W**). These results collectively suggest that the transient increases in mEPSC and spine density in 1d and 5d winners likely reflect an increase in local connectivity among VMHvl^Esr1^ cells.

**Figure 6.**
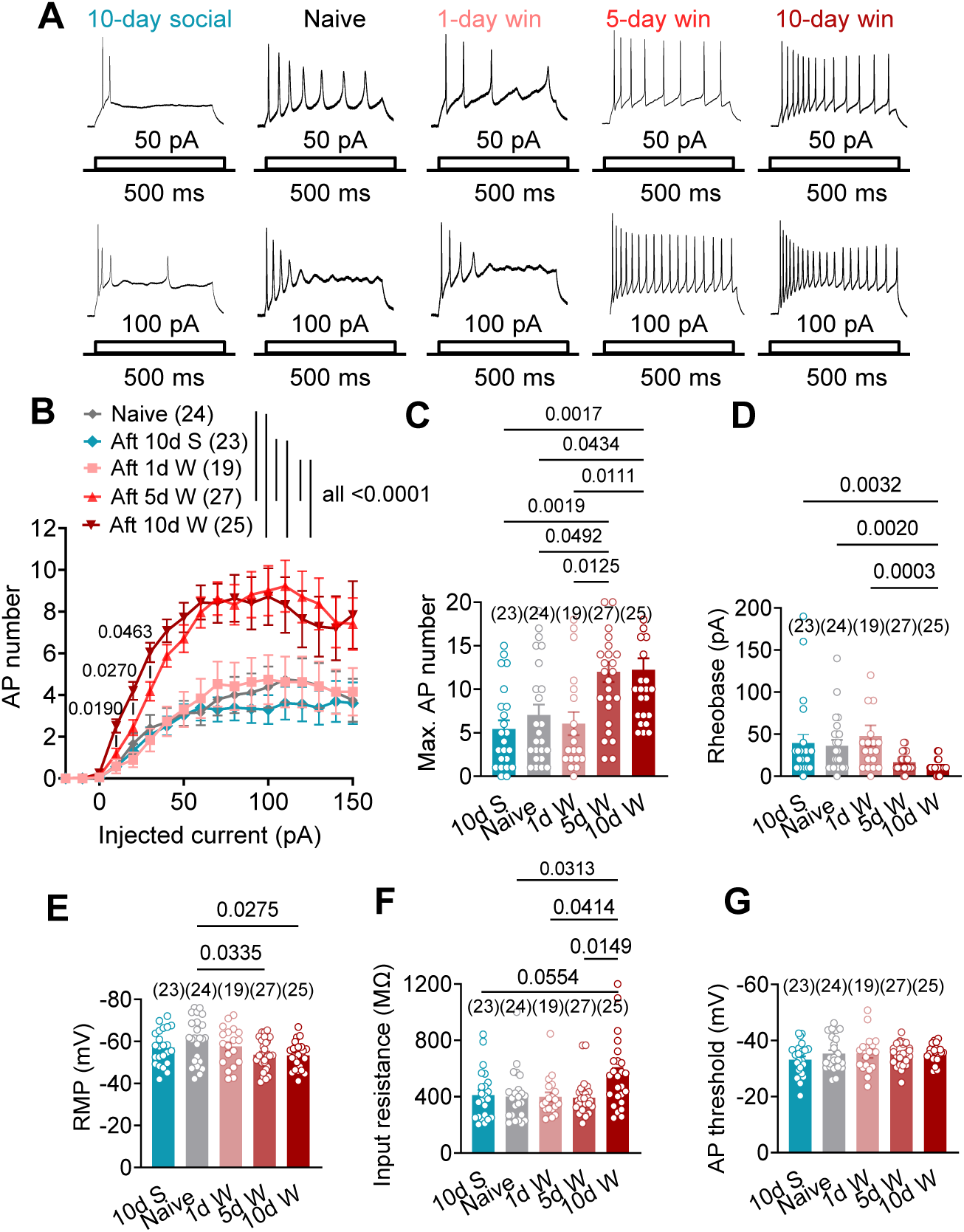
Increase of VMHvl^Esr1^ cell excitability after repeated winning. (A) Representative current-clamp recording traces of VMHvl^Esr1^ cells with 50pA and 100pA current injections. (B) Frequency-current (F-I) curve of VMHvl^Esr1^ cells in naïve mice, 10d social mice, and mice with different days of winning experiences. (C) Maximal numbers of action potentials across current steps of various groups. (D-G) Rheobase (D), resting membrane potential (E), input resistance (F), and action potential (AP) threshold (G) of VMHvl^Esr1^ cells in naïve mice, 10-day social mice, and mice with 1, 5, and 10 days of winning experiences. Bar and error bar: mean ± SEM. Circles represent individual cells. Numbers in parenthesis indicate the number of recorded cells. (B) Two-way ANOVA with Tukey’s multiple comparisons test and two-way ANOVA with repeated measures followed by Tukey’s multiple comparisons test. (C, D, G) Kruskal-Wallis test with Dunn’s multiple comparisons test. (E, F) One-way ANOVA followed by Tukey’s multiple comparisons test. All p :5 0.05 are specified. If not indicated, p > 0.05. See Supplementary Table 2 for detailed statistics.

### Changes in intrinsic properties of VMHvl^Esr1^ cells after repeated winning

We further asked whether winning experiences alter the intrinsic biophysical properties of VMHvl^Esr1^ cells using in vitro slice recording. We first constructed the spike frequency-current (F-I) curve by injecting a series of current steps (-20 pA to 150 pA) into the VMHvl^Esr1^ cells (**Figure 6A**). F-I curves in naïve and 1d winners are similar, whereas cells in 5d and 10d winners showed higher firing rates in nearly all current steps and had significantly higher maximum firing rates compared to naïve and 1d winner groups (**Figure 6A-C**). Between 5d and 10d winners, the F-I curve was steeper in 10d winners at low current steps (10-30pA), indicating increased sensitivity to small excitatory inputs (**Figure 6B**). Additionally, VMHvl^Esr1^ cells of 10d winners showed significantly lower rheobase (the minimum current to evoke a spike) than other groups; a trend of decrease was also observed in 5d winners (**Figure 6D**). Resting membrane potential (RMP) became gradually more depolarized over winning: 5d and 10d winners showed significantly depolarized RMP than naïve animals (**Figure 6E**). Interestingly, input resistance only increased in 10d, but not 1d or 5d, winners (**Figure 6F**). Lastly, the spiking threshold did not change in any groups (**Figure 6G**). None of the measured parameters in 10d social animals differed from naïve animals, suggesting that these intrinsic properties changes are winning-dependent (**Figure 6A-G**). Thus, the VMHvl cells show increased excitability after long but not short-term winning.

### The VMHvl synaptic and intrinsic plasticity are causally linked

Our results revealed three types of plasticity in the VMHvl over 10 days of repeated winning: continued potentiation of glutamatergic synapses with long-range inputs, transient increase of local connections, and delayed increase of cell excitability. We wondered whether these events are linked and what might be their trigger. Among the three types of plasticity, the PA-VMHvl potentiation occurs most rapidly. Indeed, immediately after the 10-minute RI tests, we can detect an increase in PA terminal activation-evoked LFP at the VMHvl (Figure S5A-E). This increase is likely due to Hebbian plasticity, as PA and VMHvl cells show simultaneous activation during inter-male aggression (Guo et al., 2023; Yamaguchi et al., 2020). Thus, we decided to test whether the PA-VMHvl potentiation, in the absence of winning, is sufficient to trigger the series of plasticity events, including the transient sEPSC frequency increase, spine density and morphology changes, and delayed increase in excitability.

We first confirmed that VMHvl and PA-VMHvl coactivation is sufficient to induce PA-VMHvl potentiation as described previously (Stagkourakis et al., 2020). Specifically, we expressed Chronos in the PA^Esr1^ cells and ChrimsonR in VMHvl^Esr1^ cells and implanted an optrode in the VMHvl (**Figure S5F**). We then delivered 530 nm light (25s on, 5s off, 3 times) to coactivate the PA-VMHvl terminal and VMHvl cell body and probed the PA terminal activation-evoked LFP at the VMHvl immediately before and after the light train (**Figure S5G**). We found that the 20Hz coactivation protocol (referred to as the long-term potentiation (LTP) protocol) can rapidly increase the oLFP at the VMHvl (**Figure S5H-J**). Furthermore, repeated co-stimulation of PA-VMHvl terminal and VMHvl cells over 10 days (once a day) caused an accumulated increase of oLFP, indicating a gradual strengthening of the PA-VMHvl pathway as observed over 10-day winning (**Figures 4Q-S and S5O-Q**). The pre- and post-coactivation-induced acute synaptic potentiation was also observed in vitro at the single-cell level using patch clamp recording (**Figure S5R-V**).

We then bilaterally expressed Chronos in the PA^Esr1^ cells and ChrimsonR in VMHvl^Esr1^ cells and coactivated PA-VMHvl terminal and VMHvl cells on one side of the brain daily using the LTP protocol for 1, 5, and 10 days. The other side of the brain was left unstimulated to serve as a within-animal control (**Figure 7A)**. The day after the last co-stimulation, we performed in vitro patch clamp recording of VMHvl^Esr1^ cells and measured sEPSC and cell excitability from both stimulated and control sides (**Figure 7B**). After 1-day of co-stimulation, the sEPSC frequency of the VMHvl^Esr1^ cells at the stimulated side was significantly higher than that of the control side (**Figure 7D1-E1**), while the excitability of cells of the two sides was similar (**Figure 7G1-H1**). After 5 days of co-stimulation, sEPSC frequency as well as cell excitability became significantly higher on the stimulated side than the control side (**Figure 7D2-E2, G2-H2).** Remarkably, after 10-day co-simulation, the sEPSC frequency of the stimulated side decreased to a similarly low level as the control side, while the cell excitability was significantly higher on the stimulated side than the control side (**Figure 7D3-E3, G3-H3**). No change in sEPSC amplitude was observed between the stimulation side and control side regardless of the duration of stimulation, which is consistent with a lack of sESPC amplitude change after winning (**Figure 7F)**. Similar to the intrinsic property changes after long, but not short, period of winning, we observed lower rheobase and elevated RMP of VMHvl^Esr1^ cells on the stimulated side compared to the control side after 5 and 10 days of co-stimulation, not after 1 day of co-stimulation **(Figures S6A, B, D, E, G and H)**. Furthermore, an increase in input resistance was only observed after 10-day co-stimulation, which matches the time course of input resistance change over the course of winning precisely **(Figure S6C, F, I)**. In mice injected with the GFP virus into the PA and mCherry into the VMHvl, the cell excitability in the stimulated side did not differ from the control side after 10 days of light delivery, suggesting that the light alone does not cause the plasticity (**Figure S7A-E)**.

**Figure 7.**
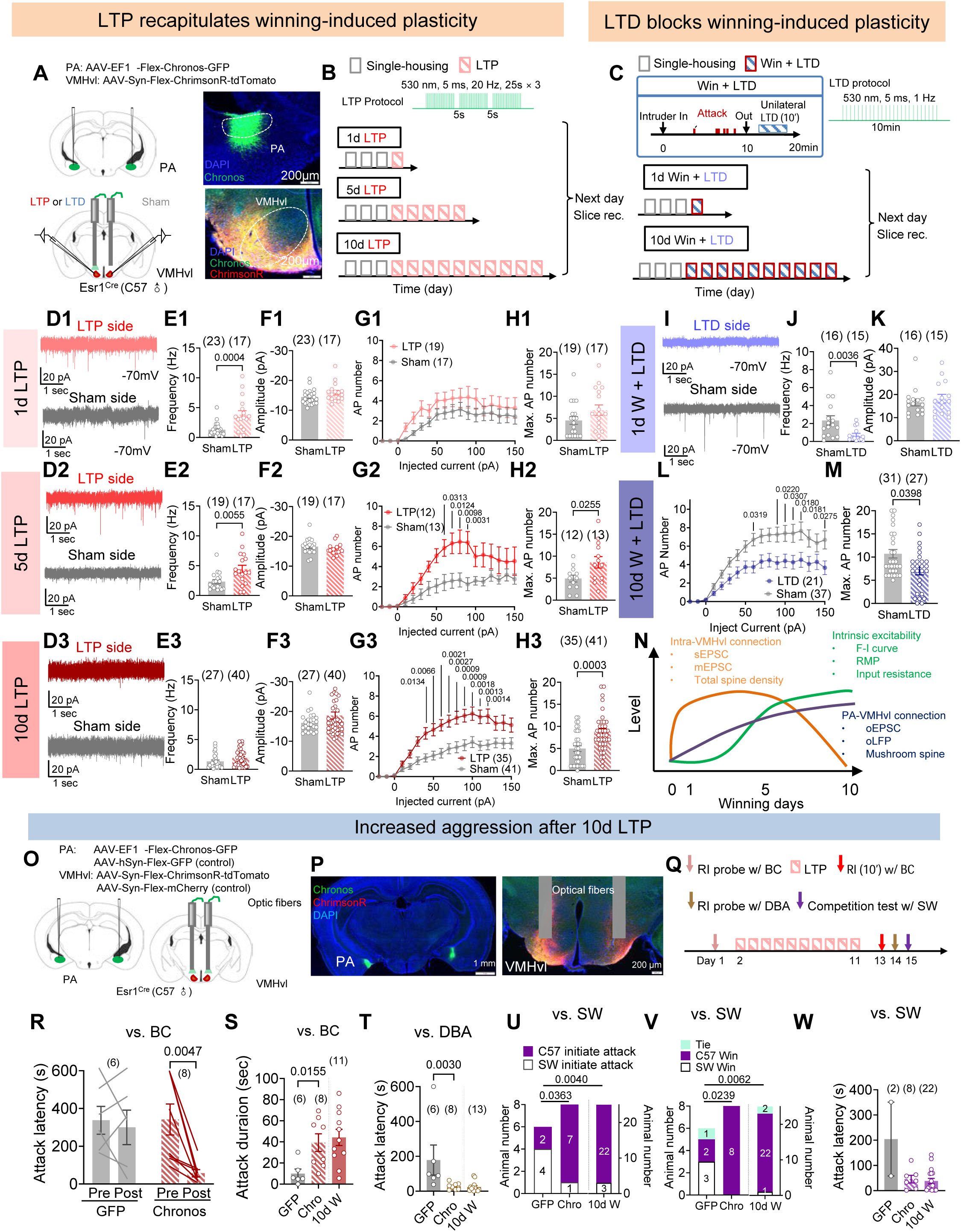
The cascade of plasticity events is causally linked and sufficient to induce the Winner effect. (A) Schematic of virus injection, light delivery and representative histology images. (B) Experimental timeline of slice recordings after LTP induction and the LTP protocol. (C) Experimental timeline of slice recordings after 1d or 10d winning paired with LTD induction and the LTD protocol. (D) Representative voltage-clamp recording traces of VMHvl^Esr1^ cells recorded from sham and 1d (D1), 5d (D2), and 10d (D3) LTP induction side. (E-F) The sEPSC frequency (E) and amplitude (F) of VMHvl^Esr1^ cells recorded from sham and 1d (E1, F1), 5d (E2, F2), and 10d (E3, F3) LTP induction side. (G) F-I curve of VMHvl^Esr^1 cells recorded from sham and 1d (G1), 5d (G2), and 10d (G3) LTP induction side. (H) The maximal number of action potentials across current steps of VMHvl^Esr1^ cells recorded from sham and 1d (H1), 5d (H2), and 10d (H3) LTP induction side. (I) Representative voltage-clamp recording traces of VMHvl^Esr1^ cells recorded from sham and LTD induction side of mice with 1 day of winning experience. (J-K) The sEPSC frequency (J) and amplitude (K) of cells on the sham and LTD sides of mice with 1 day of winning experience. (L) F-I curve of cells recorded on the sham and LTD induction sides in mice with 10 days of winning experience. (M) Maximal number of action potentials of VMHvl^Esr1^ cells recorded on the sham and LTD sides in mice with 10 days of winning experience. (N) Illustration of different types of plasticity events that occurred over the course of repeated winning. (O) Virus injection and light delivery strategy for in vivo LTP induction in freely moving mice. (P) Histology images showing virus expression and optic fiber locations. (Q) Experimental timeline for testing aggressive behaviors using different types of intruders before and after 10d LTP induction. (R) The latency to attack BC male intruders before and after 10d LTP induction. (S) The duration of attacking BC intruders after 10 days of LTP induction vs. 10 days of winning. (T) The latency to attack DBA male intruders during the probing test after 10 days of LTP induction or 10 days of winning over BC intruders. (U) The number of C57 and SW mice that initiate the 1st attack during the competition test after 10d LTP induction or 10d winning over BC intruders. (V) The results of competition tests after 10 days of LTP induction or 10 days of winning over BC. (W) The latency to attack SW male intruders after 10 days of LTP induction or 10 days of winning over BC. Bar and error bar: mean ± SEM. Circles in E, F, H, J, K and M represent individual cells. Circles and lines in R, S, T, U, V, W represent individual animals. Numbers in parenthesis in E, F, G, H, J, K, L and M indicate the number of recorded cells. Numbers in parenthesis or inside of the bars in R-W indicate the number of subject animals. (E1, E2, F1, H1, H2, S) Unpaired t test. (E3, F2, F3, H3, J, K, M, T) Mann-Whitney test. (G1, G2, G3, L, R) Two-way ANOVA with repeated measures followed by Sidak’s multiple comparisons test. (U, V) Chi-square test. All statistical tests are two-tailed. All p :5 0.05 are specified. If not indicated, p > 0.05. See Supplementary Table 2 for detailed statistics. See also Figs S5, S6, S7, S8 and S9.

The morphological change after 10-day VMHvl and PV-VMHvl coactivation also resembles that after 10-day winning. While the total spine density of VMHvl^Esr1^ cells on the stimulated side did not differ significantly from the control side, the spine composition differed drastically (**Figure S8A-C**). The proportion of mushroom spines was 40% on the stimulated side, much higher than the 11% on the control side, due to increased mushroom spine density on the stimulated side (**Figure S8B, G**). Additionally, the mushroom spine head size was significantly larger on the stimulated side than on the control side (**Figure S8H**). In contrast, the density of thin spines, filopodia, and stubby spines did not differ between stimulated and control sides (**Figure S8D-F)**.

To understand whether the PA-VMHvl pathway potentiation is necessary for the synaptic and cellular changes induced by winning, we blocked post-winning PA-VMHvl synaptic potentiation using a 1-Hz co-stimulation protocol (refer as long-term depression (LTD) protocol) as previously described (Stagkourakis et al., 2020). Specifically, immediately after the 10-min RI test during which the test animal won, we delivered 530 nm light to one side of the VMHvl at a low frequency (1Hz, 1ms) for 600 s while leaving the other side unstimulated (**Figure 7A, C**). This LTD protocol effectively weakened PA-VMHvl synaptic connection in vivo and in vitro (**Figure S5K-N, W-Z**). After applying unilateral LTD light-stimulation protocol after winning for one day, we found that the sEPSC frequency of cells on the control side was significantly higher than those on the LTD side (**Figure 7I-J**). The sEPSC amplitude did not differ between the two sides, which is consistent with the fact that winning causes no change in sEPSC amplitude (**Figure 7K**). The morphological change corroborates with the sEPSC frequency change. The total spine density increased drastically on the control side, as expected from the 1d winner, while it remained low on the LTD side (**Figure S8I-K**). The densities of all types of spines were lower on the LTD side than on the control side (**Figure S8L-O**). There is no significant difference between the mushroom spine head volume between the two sides (**Figure S8P**).

In a separate cohort animal, we performed the post-winning LTD-induction stimulation for 10 consecutive winning days and recorded the cells from the simulated and control sides the day after the last win (**Figure 7C**). While cells in the control side showed high excitability, as expected from 10d winners, the LTD side showed low excitability, comparable to that of naïve animals (**Figures 6B, 7L-M)**. Control animals injected with GFP and mCherry viruses showed increased cell excitability in both light-delivered and control sides (**Figure S7A, F-I**). These results suggest that PA-VMHvl synaptic potentiation is a prerequisite for the winning-induced sEPSC increase, spine growth, and excitability changes.

### PA-VMHvl synaptic potentiation recapitulates the winning-induced behavior changes

Given that VMHvl and PA-VMHvl coactivation can induce synaptic and cellular plasticity in the VMHvl as winning, we next asked whether the LTP protocol is sufficient to induce the winner effect behaviorally. We expressed Chronos (GFP for control) in the PA^Esr1^ cells and ChrimsonR (mCherry for control) in the VMHvl^Esr1^ cells and implanted optic fibers bilaterally above the VMHvl (**Figure 7O, P**). Four weeks after the surgery, we optogenetically induced the potentiation of the PA-VMHvl pathway bilaterally in the naïve male mice for 10 days (**Figure 7Q)**. To test the animal’s initial aggressiveness, we probed the test and control animals’ behavior towards BC intruders one day before the first day of stimulation. Then, after completing the 10-day LTP induction, we examined the behaviors towards BC, DBA, and SW intruders, each on a separate day (**Figure 7Q)**. We found that the attack latency towards BC significantly decreased while the attack duration increased after 10-day light stimulation in the test group but not in the control group (**Figure 7R, S)**. The BC-directed attack duration after 10-day co-stimulation of the test mice was comparable to that of 10-day winners (**Figure 7S).** Additionally, after 10-day co-stimulation, the test mice, but not the control mice, attacked an unfamiliar DBA mouse as quickly as the 10-day winners (**Figure 7T)**. Most strikingly, 7/8 test mice initiated the 1^st^ attack towards the strong SW mice, and all 8 won the fight while only 2/6 control mice did so (**Figure 7U, V**). Overall, the SW-directed aggression after 10-day PA-VMHvl LTP induction is comparable to that of 10d winners (**Figure 7U-W**).

Given that MUP and T levels increase after 10-day winning (**Figure 1R, S**) and their suggested causal role in the winner effect (Oyegbile and Marler, 2005), we wondered whether the 10d LTP stimulation also induces endocrine changes, which in turn contributes to the aggression increase. We thus collected the tail blood and urine samples before and after the 10-day co-stimulation and found that post-stimulation T and MUP levels did not differ from pre-stimulation levels in either test or control groups (**Figure S9A-C)**. Furthermore, we asked whether the co-stimulation changes aggression-unrelated behaviors, e.g., an increased tendency to adventure as the case after repeated winning (**Figure S2A-I**). We found that the animal’s performance in the light-dark box test did not change: the total time spent and distance traveled in the light box were similar before and after the 10-day light stimulation in test and control animals (**Figure S9A, D-E)**. These data suggest that the cascade of plasticity occurring at the VMHvl is sufficient to support winning-induced aggression increase independent of testosterone and MUP changes. The neural circuits beyond the VMHvl likely support changes in non-aggressive behaviors after repeated winning.

## Discussion

Here, we characterized changes in aggressive behaviors over repeated winning and investigated the neuroplasticity underlying these changes. We found that as male mice gain winning experiences, they shift from low and specific aggression to high and generalized aggression, and the winning probability against a stronger opponent increases. We further uncovered three forms of plasticity in the VMHvl over repeated winning, each with a distinct time course (**Figure 7N**). These neuroplasticity events collectively allow aggression to vary widely across individuals based on the fighting history.

### The winner effect

From an ethological point of view, every fight is an opportunity for an animal to assess its resource-holding potential (RHP) within its species (Parker, 1974). Winning one fight signals the relatively high RHP of that individual within the pair. Repeated winning indicates an absolute high RHP among all individuals of the species, hence encouraging the winner to engage in future agonistic encounters more readily as the risk is presumably low. Indeed, we found that after one time winning, the winning animal increases its attack specifically towards the same type of intruder that it previously defeated. As the animals gain repeated winning experiences, they show high aggression towards various intruders, even those with apparent physical superiority. In our study, however, we did not find 10d winners attack female intruders, which is considered pathological as it reduces reproductive success. Thus, the behaviors of the 10-day winners are likely still within the normal range of male mouse aggression.

We noticed that aggression level, measured by the total attack duration, towards repeatedly encountered BC intruders tends to be lower in 10d winners compared to 5d winners. This phenomenon has been reported previously: as the animal encounters the same type of intruder repeatedly, the overt attack is replaced by other dominant behaviors, e.g., lateral threat or aggressive grooming (Kudryavtseva et al., 2014). Similarly, when a group of unfamiliar male mice are housed together, the fighting frequency is the highest during the first day or two and decreases once the social hierarchy is established (Williamson et al., 2016). Thus, the decreased attack towards a repeatedly encountered weaker intruder could be considered an adaptive behavior to reduce unnecessary fighting once the winner is clear. Importantly, this decreased attack duration towards familiar intruders should not be considered a sign of reduced aggression. When the 10d winner encounters an unfamiliar intruder, especially one with a higher RHP, the aggression level is clearly higher than that of 1d or 5d winners.

The experienced winner has a higher probability of winning against the stronger opponent. While we initially attributed this increase to increased attack efficiency and precision, our results in 10-day VMHvl and PA-VMHvl co-simulated animals suggest that aggressive motivation may dominate the winning probability in the short RI tests. Previous in vivo recordings reveal that VMHvl cells are activated during aggression-seeking and attack initiation but do not vary with moment-to-moment attack movements (Falkner et al., 2016; Mei et al., 2023a). Thus, changes in VMHvl are expected to mainly affect readiness and persistence to attack but not alter the motor performance of attack.

Beyond aggression, repeated winning also enhances general adventurousness. 10d winners are more willing to explore risky environments, e.g., the light side of a light-dark box. The VMHvl does not mediate these changes, as our LTP-induced animals showed no such behavioral change. This is unsurprising, given that VMHvl cells are not activated under non-social contexts. How repeated winning affects the neural circuit beyond aggression-generation regions is an interesting topic for future studies.

### The neuroplasticity in the VMHvl over repeated winning

During winning, PA and VMHvl cells are simultaneously activated, which triggers a series of changes in VMHvl cells. First, within minutes of winning, the PA-VMHvl excitatory synapses are potentiated, revealed by increased PA-VMHvl terminal stimulation-evoked LFP at the VMHvl. Similar PA-VMHvl potentiation can be induced by simply coactivating PA-VMHvl terminals and VMHvl cells for less than two minutes, supporting the pre- and post-synaptic coactivation as the key trigger of plasticity. The winning-induced PA-VMHvl potentiation is relatively stable as the effect remains 24 hours after winning. Over 10 days of repeated winning, the PA-VMHvl connection is continuously and monotonically strengthened due to combined changes at presynaptic, e.g., increased releasing probability, and post-synaptic sites, e.g., increased AMPAR density. In 10d winners, the number of mushroom spines increases significantly. Although not directly proven, some of these newly emerged mushroom spines likely partner with PA axons as artificial PA-VMHvl pathway potentiation similarly increases the mushroom spine density, whereas PA-VMHvl depression prevents the mushroom spine from growing after repeated winning. It is worth noting that PA represents the primary extra-hypothalamic inputs to the VMHvl. Other long-range inputs, including those from BNST, MeA, and LSv, are mainly GABAergic (Yamaguchi et al., 2020). The other major source of glutamatergic inputs to the VMHvl arises from the PMv, a hypothalamic region just posterior to the VMHvl (Lo et al., 2019; Soden et al., 2016). PMv also promotes aggression and is activated by male chemosensory cues (Chen et al., 2020; Soden et al., 2016; Stagkourakis et al., 2018). Thus, the PMv to VMHvl connection may also be strengthened over repeated winning, a possibility to be investigated in future studies.

Surprisingly, PA-VMHvl and VMHvl coactivation also trigger plasticity unrelated to the PA-VMHvl pathway. One day after PA-VMHvl and VMHvl coactivation, we observed a striking increase in sEPSC frequency and VMHvl cell spine density, similar to after 1d winning. Most of these newly formed spines are thin spines and filopodia and disappear by the 10^th^ coactivation day, again as what happens after 10d winning. Two pieces of evidence suggest that these transient spines are likely formed among VMHvl cells. First, chemogenetic inhibition of VMHvl cells, which suppresses presynaptic vesicle release specifically from VMHvl cells, reduces VMHvl cell sEPSC frequency in 1d and 5d winners but not naïve animals or 10d winners. Second, brief optogenetic activation of VMHvl cells elicits more qEPSCs from VMHvl cells in 1d and 5d winners than naïve mice and 10d winners. The functional importance of the strengthened local connection remains unclear as effective tools to block VMHvl local synaptic transmission without simultaneously blocking cell body activation are currently lacking. We speculate that the increased recurrent excitation may amplify the input, synchronize the output, or maintain the spontaneous activity of VMHvl cells once triggered. Indeed, in vivo single-unit recording and miniscope imaging revealed elevated baseline activity of VMHvl cells between attack episodes and after removal of the male intruder (Lin et al., 2011; Nair et al., 2023). The mechanisms causing the local VMHvl connection to diminish in highly experienced winners remain unclear. We speculate that this may reflect homeostatic plasticity to prevent runaway excitation of the VMHvl once the cell excitability increases sufficiently.

Repeated winning also causes a delayed increase in VMHvl cell excitability. PA-VMHvl and VMHvl coactivation for 10 days mimics the winning-induced excitability increase while blocking the PA-VMHvl potentiation after daily winning prevents the excitability change, suggesting these two events, synaptic potentiation and intrinsic excitability increase, are causally linked. Indeed, studies in the last decade revealed experience-dependent changes in cell excitability in the cerebellum, hippocampus, and cortex. In all cases, NMDA receptor activation and Ca^2+^ influx are found indispensable for the “Hebbian intrinsic excitability” (Aizenman and Linden, 2000; Armano et al., 2000; Debanne et al., 2019; Zhang and Linden, 2003). In our study, the fact that synaptic potentiation occurs sooner than excitability increases suggests that a higher or longer-lasting Ca^2+^ increase is needed for intrinsic property changes compared to synaptic changes. Intriguingly, although thin spines contain fewer AMPA receptors than mushroom spines, the number of NMDA receptors across spines is less heterogeneous (Matsuzaki et al., 2001). The transiently appearing local thin spines may be a prerequisite to increased excitability by elevating and maintaining a high intracellular Ca^2+^ level.

Although the T level is elevated after repeated winning, we found that a T increase is not required for increasing VMHvl cell excitability or aggression. Repeated coactivation of PA-VMHvl and VMHvl cells can increase VMHvl cell excitability as well as inter-male aggression without altering the T level. Hence, in contrast to the popular belief (Batrinos, 2012; Oyegbile and Marler, 2005)(but see (Demas et al., 2023) for examples of T-independent aggression change), T appears to play no instructive role in enhancing aggression after winning. This does not mean that T plays no role in the winner effect. Instead, it may play a permissive role. For example, in the absence of T, e.g., through castration, VMHvl cells may decrease excitability drastically, causing the elimination of aggression altogether. Nevertheless, our data suggests that aggression increase after winning is not secondary to T increase. Instead, Hebbian synaptic and excitability plasticity is responsible for changes in the aggression circuit. We speculate that T increase may be important for non-aggression-related behavioral changes after winning, given that aggression-unrelated circuits should not be activated during winning, making Hebbian plasticity unlikely. Consistent with this possibility, repeated coactivation of PA-VMHvl and VMHvl cells that cause no T change also does not alter exploratory behaviors in non-social contexts.

### Neuroplasticity beyond the VMHvl that supports the winner effect

Our study revealed diverse forms of plasticity employed by the VMHvl cells to change its input-output relationship, including changes in long-range connection, local connectivity, and cell excitability. This multi-form plasticity enables a wide dynamic range for VMHvl operation, hence, the aggression level of an animal. Indeed, aggression is unique among innate social behaviors in its high variability across time and individuals. The fact that aggression can continuously increase over 10 days of repeated winning illustrates the wide output range of the aggression circuit. One key question is whether VMHvl is a unique site of plasticity after winning or one of the many in the aggression circuit. While answers to this question require additional experimentation, we predict the latter is likely the case. So far, over 10 brain regions have been found to modulate aggression, including MPN, VMHvl, PMv, DMH, AHN, PA, MeA, BNSTpr, LS, SuBv, VTA and PAG (Lischinsky and Lin, 2020). This list is likely to continue to grow. Recently, we simultaneously recorded 13 limbic regions in male mice and revealed synchronized activation across many sites during attack (Guo et al., 2023). Thus, Hebbian plasticity is likely to occur not only between the PA and VMHvl connection but also in other excitatory pathways. Indeed, PA also projects densely to MeA, BNSTpr, and PMv, which are all functionally important for generating aggression. PMv is a site of particular interest as it is highly glutamatergic, like the VMHvl. Stagkourakis et al. compared the PMv cell properties between aggressive (3d winning) and non-aggressive male mice and found that PMv cells in aggressive mice show higher excitability and denser intra-PMv cell connectivity (Stagkourakis et al., 2018). Future studies should address whether the VMHvl plasticity rules apply to the PMv and other aggression-related regions.

Despite being innate, the readiness to express aggression could vary hugely across individuals or within the same individual over time. Our study uncovered diverse forms of plasticity in the VMHvl over the course of winning that enhance the readiness to attack collectively. As the winning experience accumulates, the form of plasticity becomes increasingly stable, making the high aggressiveness of experienced winners a stable trait independent of fighting itself. Such biophysical coding of individuality is likely a general principle beyond aggression (Ammari et al., 2023; Mei et al., 2023b).

## Methods

### Mice

All procedures were approved by the NYULMC Institutional Animal Care and Use Committee (IACUC) in compliance with the NIH guidelines for the care and use of laboratory animals. All mice used in this study were housed under a 12h light-dark cycle (light cycle, 10 p.m to 10 a.m.), with food and water ad libitum. Room temperature was maintained between 20-22 ℃, and humidity between 30-70%, with an average of approximately 45%. Esr1^Cre^ knock-in mice with C57BL/6 background were purchased from Jackson Laboratory (Stocks #017913). The Esr1-zsGreen mouse was originally generated and kindly provided by Dr. Yong Xu at Baylor College of Medicine, and then it was bred in-house with C57 WT mice (Saito et al., 2016). Apart from the experimental animals, three strains of mice were also used as intruders in the RI tests. The group-housed BALB/c male mice (>8 weeks) were used when we aimed to provide the resident mice with winning experiences. Group-housed DBA/2 male mice (>8 weeks, Charles River) were used as unfamiliar intruders for aggression probing. Single-housed aggressive Swiss Webster male mice (>10 months, Taconic) were used as challengers to evaluate the maximum fighting ability of the test mice. In the fiber photometry recording, group-housed BALB/c male (>8 weeks) and female mice (>8 weeks) were used as stimulus animals.

### Viruses

AAV2-EF1a-Flex-Chronos-GFP (3.6 × 10^12^ vg/ml), AAV2-Syn-Flex-ChrimsonR-tdTomato (6.0 × 10^12^ vg/ml), AAV2-hSyn-Flex-GFP (3.7 × 10^12^ vg/ml) were purchased from University of North Carolina vector core. AAV2-CAG-Flex-GCaMP6f-WPRE-SV40 (2.21 × 10^13^ vg/ml) was purchased from the University of Pennsylvania vector core facility. AAV5-Syn-Chronos-GFP (≥ 5×10¹² vg/mL), AAV8-hSyn-DIO-hM4Di-mCherry (1 × 10^13^ vg/ml), AAV2-hSyn-DIO-mCherry (4 × 10^12^ vg/ml), and AAV9-Syn-ChrimsonR-tdTomato (≥ 1×10¹³ vg/mL) were purchased from Addgene. All viruses were aliquoted and stored at -80 ℃ until use.

### Stereotaxic surgery

Mice were anesthetized with 1.5% isoflurane and placed in a stereotaxic apparatus (Kopf Instruments). Viruses were injected into the targeted brain regions using glass capillaries and a nanoinjector (World Precision Instruments, Nanoliter) at 10 nL/min.

For fiber photometry recording of VMHvl^Esr1^ cells, 75 nL of AAV2-CAG-Flex-GCaMP6f-WPRE-SV40 was injected into the VMHvl (AP: -1.58 mm, ML: 0.775 mm, DV: 5.65 mm) of Esr1-Cre male mice. A 400-µm optical-fiber assembly (Thorlabs, FR400URT, CF440) was implanted 250 µm above the injection site and secured using dental cement (C&B Metabond, S380). During surgery, a head-fixation ring was also secured on the skull. All recordings started 3∼5 weeks after the virus injection.

For slice recording experiments with Esr1-zsGreen male mice. 100 nl of AAV9-Syn-ChrimsonR-tdTomato was bilaterally injected into the PA (AP, -5.6mm; ML, 2.4mm; DV, -5.1mm). All mice were used for slice recording approximately four weeks after surgery, regardless of the length of RI tests. For slice recording of Esr1-Cre male mice in LTP and LTD induction experiments, 100 nL of 100 nl of AAV2-EF1a-Flex-Chronos-GFP was bilaterally injected into the PA (AP, -5.6mm; ML, 2.4mm; DV, -5.1mm), and 100 nL of AAV2-Syn-Flex-ChrimsonR-tdTomato was bilaterally injected into the VMHvl (AP, -4.85mm; ML, 0.776mm; DV, -5.75mm). The 200 µm optic fibers (RWD Life Science, R-FOC-L200C-39NA) were bilaterally implanted 250 µm above the injection sites after virus injection and further secured with dental cement (C&B Metabond, S380). For the control group, 100 nL of AAV2-hSyn-Flex-GFP was bilaterally injected into the PA, and 100 nL of AAV2-hSyn-DIO-mCherry was bilaterally injected into the VMHvl. The light stimulation with LTP/LTD protocols and slice recording experiments were conducted four weeks after surgery. For slice recordings of qEPSC experiments, 100 nL of AAV2-Syn-Flex-ChrimsonR-tdTomato was bilaterally injected into the PA (AP, -5.6mm; ML, 2.4mm; DV, -5.1mm), and 100 nL of AAV2-EF1a-Flex-Chronos-GFP was bilaterally injected into the VMHvl (AP, -4.85mm; ML, 0.776mm; DV, - 5.75mm) in Esr1-Cre male mice. For slice recording of sEPSC with hM4Di manipulation experiments, 100 nL of AAV8-hSyn-DIO-hM4Di-mCherry was injected into the VMHvl (AP, - 4.85mm; ML, 0.776mm; DV, -5.75mm) bilaterally in Esr1-Cre male mice. The brains were used for recording at least three weeks after surgery. For initial checking of the LTP/LTD protocol using slice recording, 100 nL AAV5-Syn-Chronos-GFP was injected into the PA in C57 wild-type male mice.

For oLFP in vivo recording experiments, we injected 100 nL of AAV2-EF1a-Flex-Chronos-GFP into the PA and implanted an optrode into the VMHvl (AP, -4.7 to -4.85mm; ML, 0.776mm; DV, -5.5mm). The recording started approximately four weeks after surgery.

For testing changes in aggression after 10-day LTP protocol, we bilaterally injected 100 nL of AAV2-EF1a-Flex-Chronos-GFP into the PA (AP, -5.6mm; ML, 2.4mm; DV, -5.1mm), and 100 nL of AAV2-Syn-Flex-ChrimsonR-tdTomato into the VMHvl (AP, -4.85mm; ML, 0.776mm; DV, - 5.75mm). The 200 µm optic fibers (RWD Life Science, R-FOC-L200C-39NA) were bilaterally implanted 250 µm above the injection sites after virus injection and further secured with dental cement (C&B Metabond, S380). For the control group, 100 nL of AAV2-hSyn-Flex-GFP was bilaterally injected into the PA, and 100 nL of AAV2-hSyn-DIO-mCherry was bilaterally injected into the VMHvl. The light stimulation with LTP protocols and behavior experiments were performed four weeks after surgery.

### Behavior tests and analysis Resident-intruder tests

The test mice were always single-housed resident mice. In the daily winning training and aggression tests, a group-housed non-aggressive adult male BALB/c mouse was introduced into the test mouse’s home cage and allowed to interact freely with the resident mouse for 10 min. Resident mice that attacked the intruders within 10 minutes were identified as “Winners.” Mice that did not attack during the 10-minute interaction period were identified as “Social animals.”

Animals that attacked the intruder for 1 day, 5 consecutive days, and 10 consecutive days constitute 1d W, 5d W, and 10d W groups, respectively. In the 10d S group, mice showed no aggressive behaviors over the 10 days of RI tests. Animals that showed unstable aggression across days were excluded. In the RI probe test, an adult male DBA mouse (group-housed) was introduced into the home cage of the test mouse and immediately removed after the resident mouse attacked the DBA mouse. In the competition test, an adult male SW mouse (single-housed) was introduced into the test mouse’s home cage and removed once the outcome was clear. The loser typically freezes in the corner, shows submissive postures, or escapes from the winner when approached. In contrast, the winner walks freely around the cage and initiates all attacks.

### Locomotion test

We measured the animal’s locomotion in its home cage and a large open field arena. In the home cage, the top-view videos were acquired using a camera (Balser, acA640-100gm) after removing the cage lid, water bottle, and food. Trajectories and distances traveled were analyzed by ANY maze software (ANY-maze). Locomotion tests in the open field were performed following the previously described method (Kraeuter et al., 2019). Each animal was placed into the center of the open field arena (40 cm x 40 cm x 35 cm) (Stoelting, #60101) illuminated at approximately 90 lux. The distance traveled over the 5 minutes was analyzed using ANY maze software.

### Light-dark box test

The light-dark test was performed using a previously reported method with minor modifications (Takao and Miyakawa, 2006). The light-dark box consists of a starting dark box (W × L × H: 40 cm × 20 cm × 35 cm) and a lightbox (W × L × H: 40 cm × 20 cm × 35 cm), separated by a divider with an opening (Stoelting, #63101). Mice were placed into the start box with the opening blocked by a plastic plate. During the test, the plate was removed to allow the mouse to move freely between the boxes for 10 minutes. The time spent and the distance traveled in the light box were recorded and analyzed using the ANY Maze software.

### Animal tracking and behavior analysis

Animal behaviors were videotaped using a top-view camera (Edmund, 89533) controlled by StreamPix (Norpix) at 25 fps. For the animal tracking and locomotion analysis in the light-dark box test, home cage, and open field test, the recorded videos were analyzed offline using ANY-maze software. For RI, aggression probing, and competition tests, “investigate” and “attack” were manually annotated frame by frame using custom software written in MATLAB (https://pdollar.github.io/toolbox/). “Investigate” is defined as nose contact with any body part of the target mouse. “Attack” is defined as a series of actions by which the male mouse lunges, bites, chases, and pushes the target mouse.

### Measure MUP and testosterone

Urine was collected and processed to determine MUP concentration as described previously (Ning et al., 2020; Zhang et al., 2020). The urine samples were stored at -20 ℃ after collection until analysis. We measured total protein and creatinine levels using QuantiChrom Protein Creatinine Ratio Assay Kit (BioAssay Systems, #DPCR-100) and calculated their ratio.

We followed a previous method to measure the testosterone in the mouse tail blood (Yoo et al., 2020). Mice were anesthetized with 1.5% isoflurane and placed in a stereotaxic apparatus. We then collected the tail blood and centrifugated it (4 ℃, 1500 RCF) for 10 minutes to isolate the serum. The serum was stored at -80 °C until all samples were collected. We then measured the testosterone levels in the serum using an ELISA Kit (Crystal Chem, #80552) based on the manufacturer’s instructions.

### Fiber photometry recording

The fiber photometry setup was as we previously described (Falkner et al., 2016; Hashikawa et al., 2017). Briefly, a 390-Hz sinusoidal blue LED light (30 µW) (LED light: M470F1; LED driver: LEDD1B; both from Thorlabs) was bandpass filtered (passing band: 472 ± 15 nm; FF02-472/30-25, Semrock) and delivered to the brain to excite GCaMP6f. The emission lights traveled back through the same optical fiber, were bandpass filtered (passing bands: 535 ± 25 nm; FF01-535/505, Semrock), passed through an adjustable zooming lens (SM1NR01, Thorlabs; Edmun optics no. 62-561), were detected by a Femtowatt Silicon Photoreceiver (Newport, 2151) and recorded using a real-time processor (RZ5, TDT). The envelope of the 390-Hz signals reflected the intensity of GCaMP6f and was extracted in real-time using a custom TDT OpenEX program. The signal was low-pass filtered with a cut-off frequency of 5 Hz.

Three to four weeks after surgery, the mice were habituated to head fixation for at least 3 days, 30 minutes a day. On the day of recording, the mice were head-fixed, and an anesthetized male or female Balb/C mouse or a toy mouse was delivered to ∼2 mm in front of the nostrils of the recording mouse using a linear track 6 times, each for 10 s and spaced by 60 s (2 minutes in between male and female mice or between the female mouse and toy mouse). The responses of the VMHvl^Esr1^ cells to sensory cues were recorded for 3 days to ensure that the signal was stable. Then, the mice underwent repeated winning or social interaction in a daily 10-minute RI test. The neural responses to various stimuli to headfixed animals were probed on the day after 1, 5, and 10 days of RI tests. The head-fixed photometry recording was carried out 4-6 hours before the RI test if they were conducted on the same day. All the recordings were done at a similar time of the day throughout the experiment.

To analyze the recording data, the MATLAB function ‘msbackadj’ with a moving window of 25% of the total recording duration was first applied to obtain the instantaneous baseline signal. The instantaneous ΔF/F value was calculated as (F_raw_-F_baseline_)/F_baseline_. The post-stimulus histograms (PSTHs) of ΔF/F aligned to the onset of each stimulus presentation were constructed for each mouse and then averaged across mice. The average ΔF/F response was calculated as the mean ΔF/F signal during the stimulus delivery period, averaged first across trials and then animals.

### oLFP recording

We followed our previously published method for optrode recording in freely moving mice (Lin et al., 2011; Wong et al., 2016). Briefly, a movable bundle containing 16 tungsten microwires (13 µm in diameter each; California Fine Wire) and a 100-µm optical fiber (RWD Life Science, R-FOC-BL100C-22NA) was implanted into the VMHvl during surgery. We also used a simplified unmovable optrode that was constructed by attaching a single tungsten electrode (MicroProbes for Life Science, WE30030.5A3) to a 100-um optic fiber. The recorded signal was amplified using a head-mounted headstage (Tucker Davis Technology, LP16CH), passed through a torqueless, feedback-controlled commutator (Tucker Davis Technology, AC32), and digitized using a commercial amplifier (RZ5, Tucker Davis Technology).

Four weeks after surgery, we connected the implanted electrode with the headstage and delivered 0.5 - 2 mW, 5 ms, 0.1 Hz, 470 nm light pulses (Shanghai Dream Lasers Technology) to probe the oLFP. The animal then underwent a 10-minute RI test. Afterward, the oLFP was again probed. The procedure was repeated for 10 days. The light intensity for probing the LFP stayed constant across days for each animal.

To determine the oLFP change after the LTP and LTD stimulation protocol, we first probed the oLFP responses using 0.5 - 2 mW, 5 ms, 0.1 Hz, 470 nm light pulses for 3-5 min. We then applied the LTP protocol (530 nm, ∼1.3 mW, 20 Hz, 5 ms, 25 s on and 5 s off for 3 times) or LTD protocol (530 nm, ∼0.13 mW, 1 Hz, 5 ms, 600 s) (Changchun new industries optoelectronics technology). Immediately after the LTP or LTD stimulation protocol was completed, we again probed the oLFP for 5-10 min. To understand whether the LTP protocol-induced oLFP change accumulates over days, we performed LTP stimulation once a day for 10 consecutive days and probed the oLFP response daily before applying the LTP protocol.

### In vivo optogenetic modification of PA-VMHvl pathway

The implanted optical fibers (RWD Life Science, R-FOC-L200C-39NA) were connected with 200 µm multimode patch cords (Thorlabs, FT200EMT) through matching sleeves (Thorlabs, ADAL1) to deliver light. Four weeks after the surgery, we habituated the animals to the head fixation and optic fiber connecting procedures several times. For behavioral experiments, the light was delivered bilaterally daily for 10 days. For slice recording, we unilaterally applied the LTP stimulation protocol (530 nm, ∼2 mW, 20 Hz, 5 ms, 25 s on, 5 s off, 3 times) for 1, 5, and 10 days. All animals were single-housed and did not encounter any intruder. To assess the effect of blocking PA-VMHvl potentiation on winning-induced plasticity, we subjected the test animal to an RI test with a BC intruder for 10 minutes. Then, for animals that won the RI tests, we applied the LTD stimulation protocol (1Hz, ∼0.5 mW, 5ms, 600s) unilaterally immediately after the RI test for 1 day or 10 consecutive days.

### Patch clamp slice electrophysiological recording

All the slice recordings were performed on the day after the final behavior test or light stimulation and, if applicable, three to four weeks after the virus injection. Mice were anesthetized with isoflurane, and the brains were submerged in the oxygenated ice-cold cutting solution containing (in mM) 110 choline chloride, 25 NaHCO_3_, 2.5 KCl, 7 MgCl_2_, 0.5 CaCl_2_, 1.25 NaH_2_PO_4_, 25 glucose, 11.6 ascorbic acid and 3.1 pyruvic acid. The coronal VMHvl brain sections (275 uM in thickness) were cut using the Leica VT1200s vibratome and collected into the oxygenated artificial cerebrospinal fluid (ACSF) solution containing (in mM) 125 NaCl, 2.5 KCl, 1.25 NaH_2_PO_4_, 25 NaHCO_3_, 1 MgCl_2_, 2 CaCl_2_ and 11 glucose at 32-34 ℃ and incubated for 30 min. Then, the sections were transferred to room temperature until use. During recording, we moved a VMHvl section into the recording chamber perfused with oxygenated ACSF and performed the whole-cell recordings. The recorded signals were acquired with MultiClamp 700B amplifier (Molecular Devices) and digitized at 20 kHz using DigiData1550B (Molecular Devices). The stimulation and recording were conducted using the Clampex 11.0 software (Axon instruments). The intracellular solution for current-clamp recording contained (in mM) 145 K-gluconate, 2 MgCl_2_, 2 Na_2_ATP, 10 HEPES, 0.2 EGTA (286 mOsm, pH 7.2). The intracellular solution for the voltage clamp recording contained (in mM) 135 CsMeSO_3_, 10 HEPES, 1 EGTA, 3.3 QX-314 (chloride salt), 4 Mg-ATP, 0.3 Na-GTP, and 8 sodium phosphocreatine (pH 7.3 adjusted with CsOH). The recorded data were analyzed using Clampfit (Molecular Devices).

Esr1 cells in the VMHvl were labeled with zsGreen and identified with an Olympus 40× water-immersion objective with a GFP filter. The cell membrane potential was held at -70 mV to record the sEPSCs, mEPSCs, and oEPSCs, and 0 mV to record sIPSCs, mIPSCs, and oIPSCs. For mEPSCs and mIPSCs recordings, we bath applied 1µM tetrodotoxin citrate (TTX, Tocris). To activate the Chronos- or ChrimsonR-expressing axons in the VMHvl, 0.1Hz, 1ms, 470nm, or 605nm light pulses (pE-300 white; CoolLED) were delivered to the recorded sections through the objective. To determine the AMPA and NMDA receptor-mediated currents, we bath applied 2 µM SR-95531 (a GABA_A_ receptor antagonist) and held the cells at -70 mV for AMPAR-mediated current recording and +40 mV for NMDAR-mediated current recording. The AMPAR-mediated current was the maximum value of the EPSC trace. The NMDAR-mediated current was calculated as the average value between 60 and 65 ms after the light onset. In the LTP and LTD induction experiments (**Figure S5R-Z**), the cells were recorded blindly within the VMHvl (without zsGreen guidance). We probed the oEPSCs using 0.1Hz, 1ms, 470nm LED light delivered through the objective for 3min (∼20 trials), and then delivered a train of light pulses (20Hz, 5ms, 25s, 5s interval, 3 times), and then probed oEPSCs for 10min after stimulation. In the LTD experiments, we delivered 1Hz, 5ms, 470nm light for 600s after 3 min probing of oEPSCs. We also recorded the oEPSCs for 10 min after light stimulation.

We performed current-clamp recordings to determine the intrinsic excitability of VMHvl^Esr1^ cells. We recorded the voltage changes of the cells in response to a series of 500ms current steps from -20 pA to 150 pA in 10 pA increments. In addition, the intrinsic properties of each cell, such as rheobase, resting potential, input resistance, and AP threshold, were characterized. The rheobase was defined as the minimum current capable of inducing the first action potential. The resting potential was measured directly after rupturing the cell membrane with 0 pA holding current. The input resistance was read from the membrane test window of the software after membrane break-in during recording (Clampex 11.0 software). The AP threshold for the spike of each neuron was calculated as the voltage at which ΔV/Δt reached 5% of the maximum AP rise slope.

For the sEPSC recordings in hM4Di-expressing mice, the cell was held at -70mV, and sEPSCs were recorded before and after 10 min 10 μM CNO perfusion. For the light-evoked qEPSC recordings, Ca^2+^ in ACSF was replaced with 4mM Sr^2+^, 1 μM TTX and 100 μM 4AP were added to the ACSF before recording. 605 nm and 470 nm light pulses (1ms, 0.1Hz) were delivered to elicit the glutamate release from PA^Esr1^ and VMHvl^Esr1^ cells, respectivley. We detected the evoked qEPSCs 0.5-1.5s after the light onset and calculated the average number of qEPSCs from 5 sweeps using Clampfit 11 (Molecular Devices).

### Dendritic spine labeling and imaging

To obtain the cell dendritic spine morphology, we added 1% biocytin (Tocris) to the internal solution of whole-cell patch clamp recording and recorded the cell for at least 15 min before withdrawing the electrode. After the recording, the sections were fixed using 4% paraformaldehyde overnight. The free-floating brain sections were washed 3 times with phosphate-buffered saline (PBS) (10 min), followed by 1 hr blocking in 0.3% NDS PBST (0.3% Triton X-100 in PBS with 10% normal donkey serum) at room temperature. The sections were then incubated with Alexa Fluor 647 conjugated streptavidin (Thermo Fisher Scientific, 1:250 dilution) in 0.3% NDS PBST at 4℃ for 72 h. The sections were washed with 1× PBS and mounted onto super-frost slides (Thermo Fisher Scientific, 12-550-15). The z-stack images of dendritic spine segments were acquired using a confocal microscope (Zeiss LSM 700 microscope) with 64× oil objective with 0.18 µm step size. The different types of spines were identified and classified using Neurolucida 360 (MBF Bioscience). Five segments were analyzed for each cell, and the averaged data were used for further comparisons across groups.

### Immunohistochemistry

Esr1-zsgreen mice were perfused transcardially with PBS, followed by 4% paraformaldehyde in PBS. The brains were extracted, post-fixed in 4% PFA for 2 ∼ 3 hours at 4 °C followed by 48 hours in 30% sucrose, and then embedded in OCT compound (Fisher Healthcare) and frozen on dry ice. 50 μm thick coronal brain sections were cut using a cryostat (model #CM3050S, Leica Biosystems) and collected in PBS. After that, the brain slices were washed with PBS (1 × 10 minutes) and blocked in PBS-T (0.3% Triton X-100 in PBS) with 5% normal donkey serum (NDS, Jackson Immuno Research) for 30 minutes at room temperature. The slices were then incubated in primary antibody diluted in blocking solution (rabbit anti-Esr1, 1:10000, Millipore, Cat# 06-935) at 4 °C for 16-20 hours, washed with PBS-T (3 × 10 minutes), incubated in secondary antibody and Nissl diluted in 5% NDS containing PBS-T (Cy3 donkey anti-rabbit IgG, 1:500, Jackson Immuno Research, Cat# 711-165-152; 435/455 Blue Fluorescent Nissl Stain, 1:200, Thermo Fisher Scientific, Cat# N21479) for 4 hours, and then washed with PBS-T (2 × 10 minutes). After drying, Slides were covered using a mounting medium (Fluoromount, Diagnostic BioSystems, #K024). Sections were imaged using a slide scanner (Olympus, VS120) and a confocal microscope (Zeiss, LSM 800).

### Statistics

No statistical methods were used to pre-determine sample sizes, but our sample sizes are similar to those reported in previous publications (Nordman et al., 2020; Stagkourakis et al., 2020; Wei et al., 2023; Yin et al., 2022). All experiments were conducted using 2-3 cohorts of animals. The results were reproducible across cohorts and combined for final analysis. Statistical analyses were performed using Prism10 (GraphPad Software). All statistical analyses were two-tailed. Parametric tests, including one sample t-test, paired t-test, unpaired t-test, and One-way ANOVA, were used if distributions passed Kolmogorov–Smirnov (for sample size ≥ 5) or Shapiro-Wilk tests (for sample size < 5) for normality or else nonparametric tests, including one sample Wilcoxon test, Wilcoxon matched-pairs signed rank test, Mann-Whitney test, and Kruskal-Wallis test were used. For comparisons across multiple groups and variables, Two-way ANOVA was used without formally testing the normality of data distribution. Following Two-way ANOVA, differences between groups were assessed using Sidak’s multiple comparison test or Tukey’s multiple comparisons test based on the Prism recommendation. When more than two one-sample t-tests were performed, the p values were adjusted using Holm-Šídák correction. All p values < 0.05 are indicated. If not specified, p>0.05. Error bars represent ± SEM. For detailed statistical results, including exact p values, F values, t values, degree of freedom, and cohort number, see Supplementary Table 2.

## Acknowledgement

This research was supported by NIH grants R01MH124927, R01MH101377 and U19NS107616 (DL).

---

**Table S1.** Characterization of different strains of mice used in the study.

| Resident Strain | Intruder Strain | Sex | Age | Hosing Condition | Body weight (mean ± STD) | Fur color |
| --- | --- | --- | --- | --- | --- | --- |
| C57BL/6 |  | Male | 2-5 months | Single-housing | 25.0 ± 3g | Black |
|  | Balb/c | Male | 2-3 months | Group-housing (5 mice/cage) | 21.0 ± 3g | White |
|  | DBA/2 | Male | 2-3 months | Group-housing (5 mice/cage) | 21.0 ± 2g | Dilute brown |
|  | Swiss Webster | Male | >10 months | Single-housing | 40.0 ± 3g | White |

**Figure S1.**
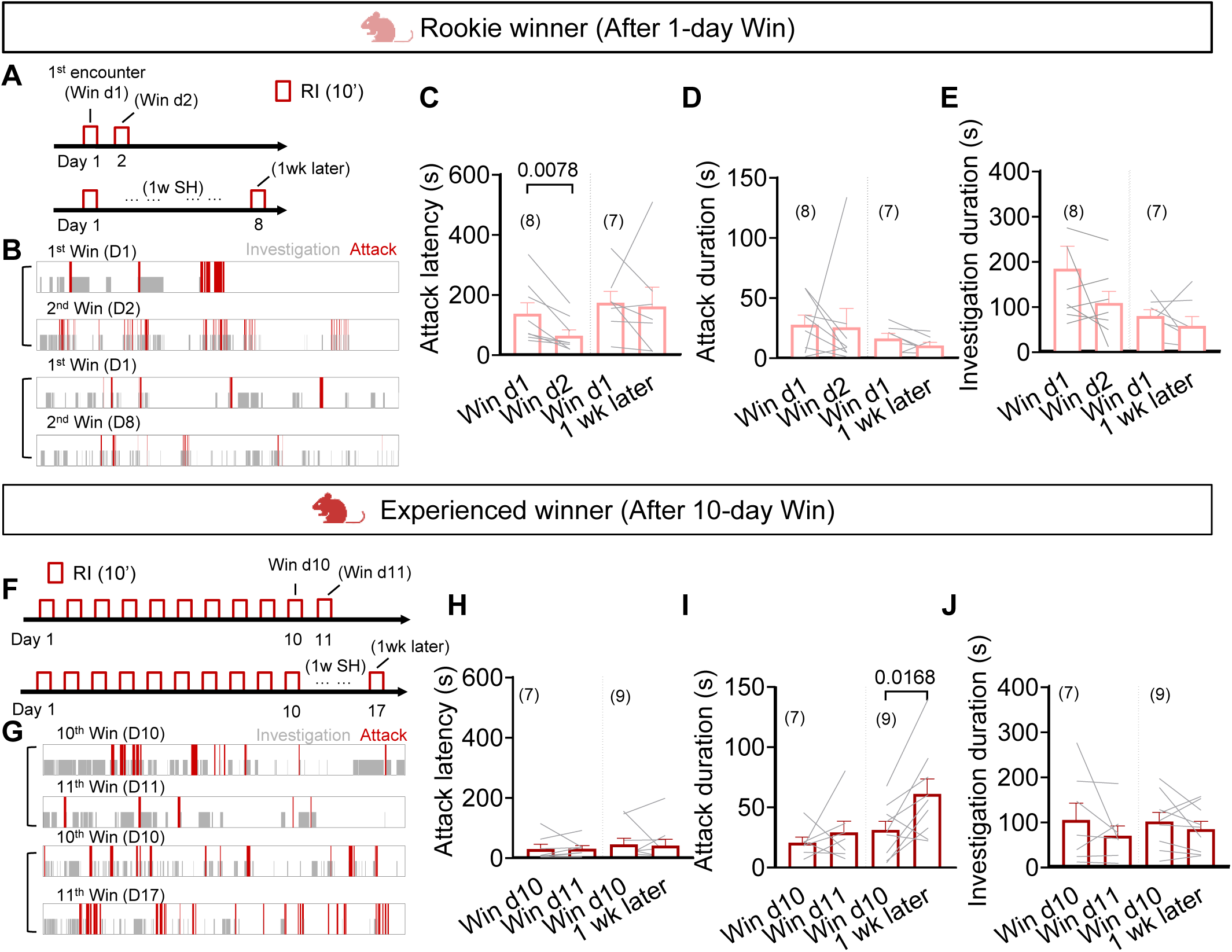
Stability of aggression in rookie and experienced winners. Related to Figure 1. (A) Behavior paradigm to test aggression levels 1 and 8 days after 1 day of winning. (B) Raster plots of attack and investigation during the RI tests 1 and 8 days after 1 day of winning. (C-E) Latency to attack (B), attack duration (C), and investigation duration (D) during the 1^st^ RI tests and 2^nd^ RI tests 1 and 8 days later. (F) Behavior paradigm to test aggression levels 1 and 8 days after 10 days of winning. (G) Raster plots of attack and investigation during the RI tests 1 and 8 days after 10 days of winning. (H-J) Latency to attack (H), attack duration (I), and investigation duration (J) during the 10^th^ RI tests and the 11^th^ RI tests 1 and 8 days later. Bars and error bars represent mean ± SEM. Lines represent individual animals. Numbers in parenthesis indicate the number of subject animals. (B (Win d1 vs. Win d2), C (Win d1 vs. Win d2), F, G (Win d10 vs. Win d11)) Wilcoxon matched-pairs signed rank test. (B (Win d1 vs. 1w later), C (Win d1 vs. 1w later), D, G (Win d10 vs. 1w later), H) Paired t test. All statistical tests are two-tailed. All p ≤ 0.05 are specified. If not indicated, p > 0.05. See Supplementary Table 2 for detailed statistics.

**Figure S2.**
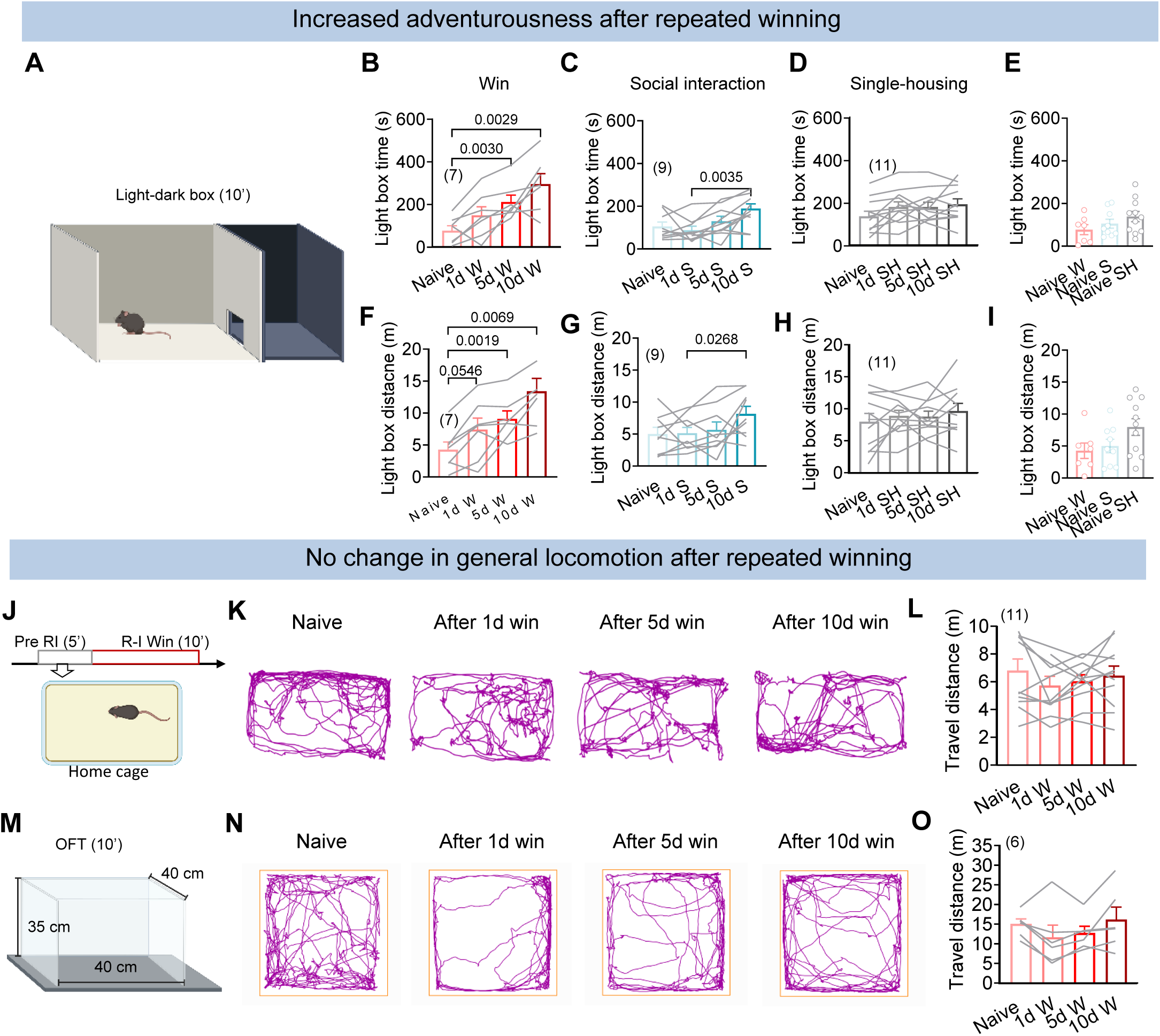
Increase in adventurousness in mice with repeated winning experiences. Related to Figure 1. (A) Schematic representation of the light-dark box. (B-D) Time spent in the light box in mice with various days of winning experiences (B), social interactions (C), and single-housing (D). (E) No difference in time spent in the light box in naïve mice before winning, social interaction, and single housing experiences. (F-H) The total travel distance in the light box with various days of winning experiences (F), social interactions (G), and single-housing (H). (I) No difference in travel distance in the light box in naïve mice before winning, social interaction and single housing experiences. (J) Measuring the locomotion in the home cage (-5-0 min before the RI test). (K) Representative tracking of the test animal’s body center in the home cage after various days of winning. (L) Travel distance in home cage in mice with various days of winning experiences. (M) Measuring the locomotion in an open field arena. (N) Representative tracking of the test animal’s body center in the open field arena after various days of winning. (O) Travel distance in the open field in mice with various days of winning experiences. Bars and error bars represent mean ± SEM. Circles and lines represent individual animals. Numbers in parenthesis indicate the number of subject animals. (B, C, D, F, G, H, O) One-way ANOVA with repeated measures followed by Tukey’s multiple comparisons test. (E, I) One-way ANOVA followed by Tukey’s multiple comparisons test. (L) Friedman test with Dunn’s multiple comparisons test. All p≤0.05 are specified. If not indicated, p> 0.05. See Supplementary Table 2 for detailed statistics.

**Figure S3.**
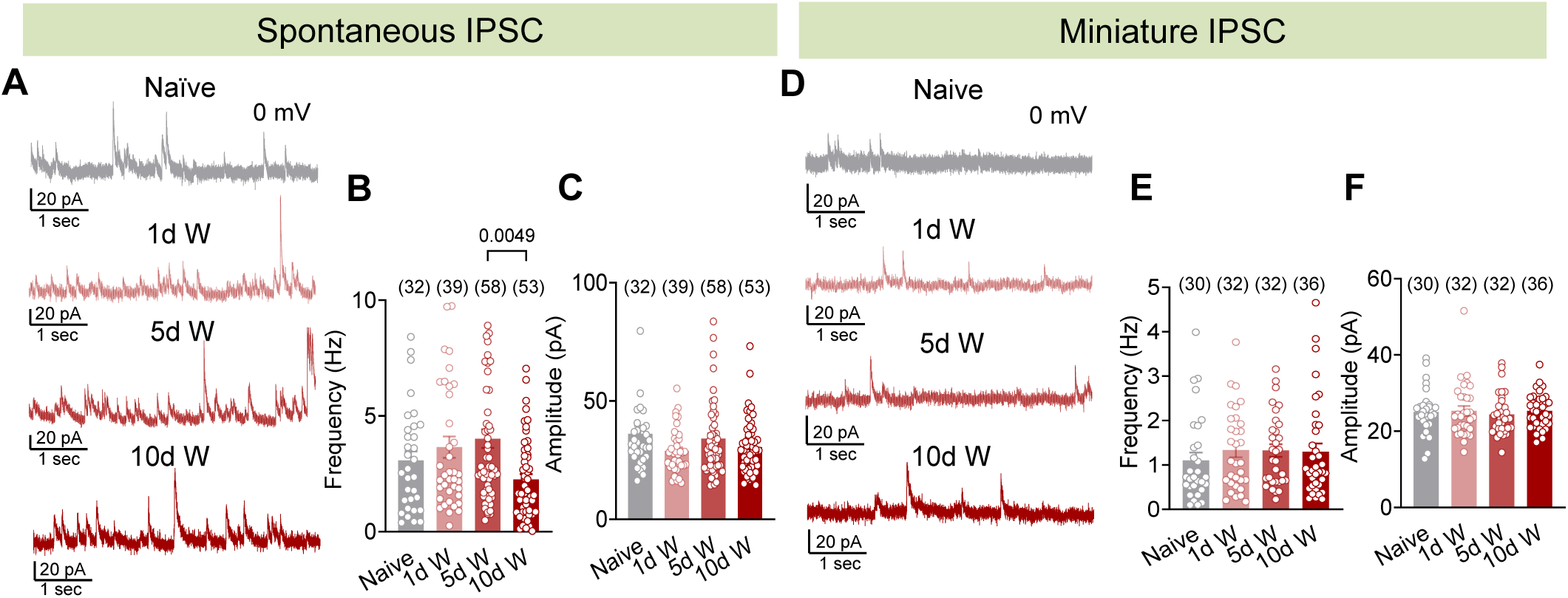
sIPSC and mIPSC in naïve mice and mice with 1d, 5d, and 10d winning. Related to Figure 3. (A) Representative voltage-clamp recording traces of VMHvl^Esr1^ cells in animals with various days of winning experiences. The holding potential is 0mV. (B, C) The sIPSCs frequency (B) and amplitude (C) of VMHvl^Esr1^ cells in animals with various days of winning experiences. (D) Representative voltage-clamp recording traces of VMHvl^Esr1^ cells in the presence of 1 µM TTX. The holding potential is 0mV. (E, F) The mIPSCs frequency (E) and amplitude (F) of VMHvl^Esr1^ cells in animals with various days of winning experiences. Bars and error bars represent mean ± SEM. Circles represent individual cells. Numbers in parenthesis indicate the number of recorded cells. (B, C, E, F) Kruskal-Wallis test with Dunn’s multiple comparisons test. All p ≤ 0.05 are specified. If not indicated, p > 0.05. See Supplementary Table 2 for detailed statistics.

**Figure S4.**
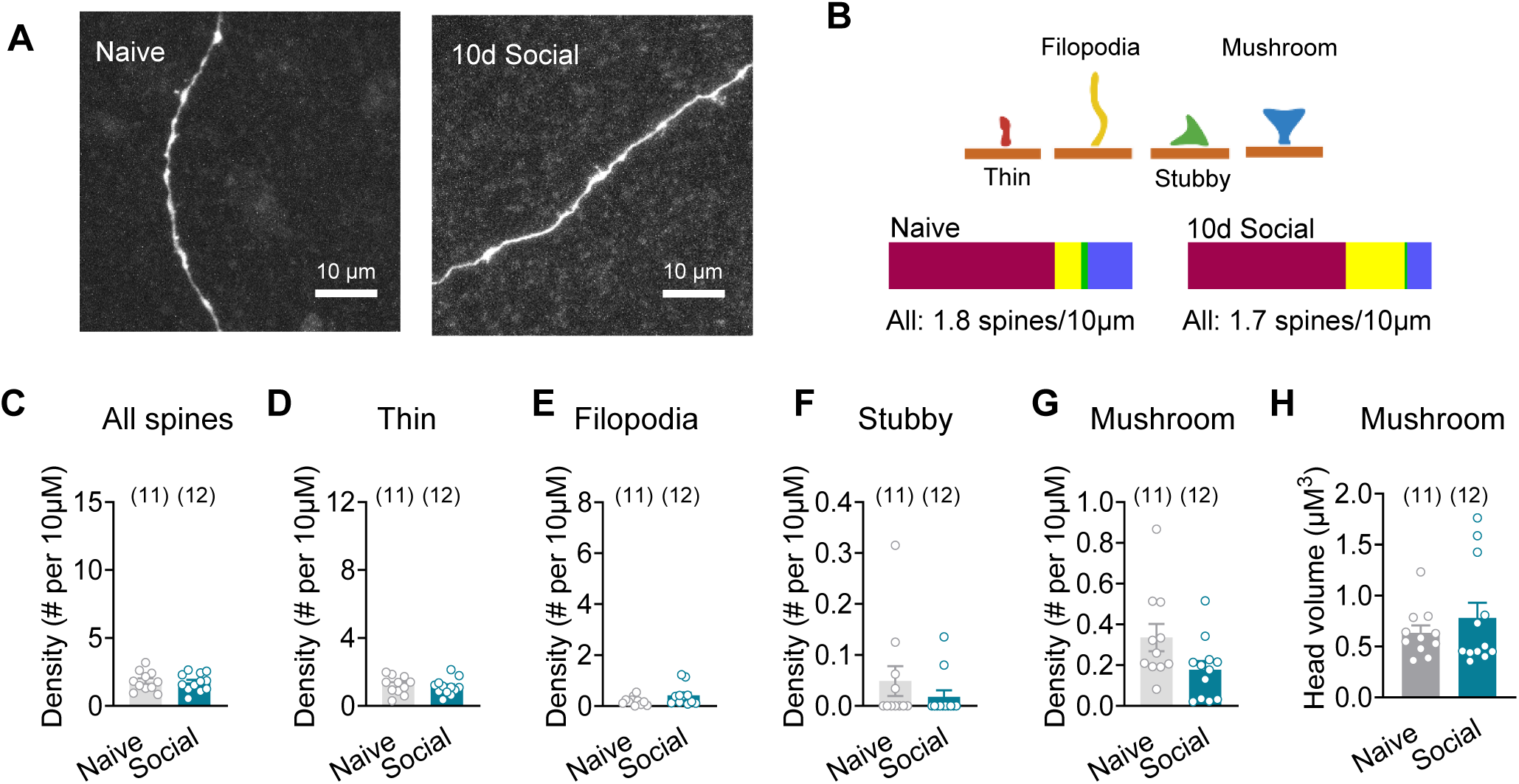
Repeated social interaction does not increase mushroom spine density. Related to Figure 3. (A) Representative images of VMHvl^Esr1^ cell dendrites in naïve (left) and 10-day social (right) groups. (B) Schematic illustration of different types of spines (top) and their proportions in naïve and 10-day social groups (bottom). (C) Total spine density in naïve and 10-day social groups. (D-G) The density of thin spines (D), filopodia (E), stubby spines (F), and mushroom spines (G) in naïve and 10- day social groups. (H) The head volume of mushroom spines in naive and 10-day social groups. Bars and error bars represent mean ± SEM. Circles represent individual cells. Numbers in parenthesis indicate the number of cells. (C, D, G) Unpaired t test. (E, F, H) Mann-Whitney test. All statistical tests are two-tailed. All p≤0.05 are specified. If not indicated, p> 0.05. See Supplementary Table 2 for detailed statistics.

**Figure S5.**
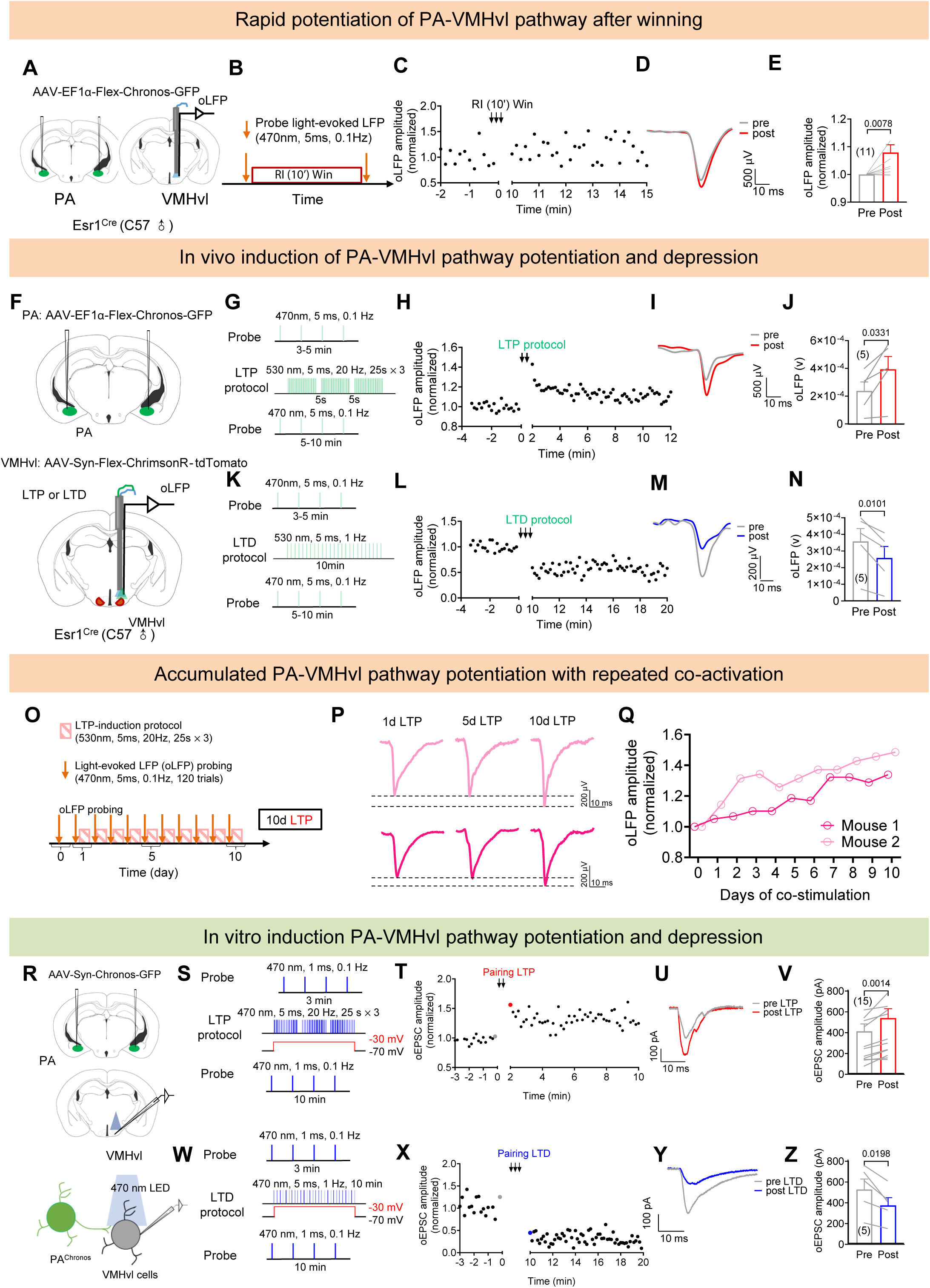
LTP and LTD stimulation protocols effectively alter PA-VMHvl connection strength. Related to Figure 7. (A) Experimental design for in vivo probing of VMHvl LFP evoked by PA terminal stimulation. (B) Experimental timeline. (C) Representative normalized oLFP before and after winning a BC male intruder in the 10-min RI test. (D) Averaged oLFP traces of a representative animal before and after winning. (E) Normalized amplitude of oLFPs before and after winning. (F) Experimental strategies for in vivo manipulation of PA-VMHvl connection strength. (G) In vivo light delivery protocol for LTP induction and oLFP probing. (H) Normalized oLFP amplitude before and after LTP induction of a representative animal. (I) Averaged traces of oLFP before and after LTP induction of a representative animal. (J) Normalized oLFP amplitude before and after LTP induction of all animals. (K) In vivo light delivery protocol for LTD induction and oLFP probing. (L) Normalized oLFP amplitude before and after LTD induction of a representative animal. (M) Averaged traces of oLFP before and after LTD induction of a representative animal. (N) Normalized oLFP amplitude before and after LTD induction of all animals. (O) Experimental timeline for 10d in vivo LTP induction. (P) Exemplar trace of oLFP after 1, 5, and 10 days of LTP induction. (Q) Normalized oLFP amplitude over 10d LTP induction. The probing occurred before the daily LTP induction (R) Experimental design for in vitro measuring of VMHvl cell EPSC evoked by PA terminal stimulation. (S) In vitro light delivery and voltage step protocol for LTP induction and oEPSC recording. (T) Normalized oEPSC amplitude before and after LTP induction of a representative cell. (U) Representative oEPSCs (the gray and red dots in T) before and after LTP induction. (V) oEPSC amplitude before and after LTP induction. The amplitude was calculated as an average of the last 5 EPSC traces before induction and the first 5 EPSC traces after induction. (W) In vitro light delivery and voltage step protocol for LTD induction and oEPSC recording. (X) Normalized oEPSC amplitude before and after LTD induction of a representative cell. (Y) Representative oEPSCs (the gray and blue dots in X) before and after LTD induction. (Z) oEPSC amplitude before and after LTD induction. The amplitude was calculated as an average of the last 5 EPSC traces before induction and the first 5 EPSC traces after induction. Bars and error bars represent mean ± SEM. Lines in E, J, N represent individual animals. Lines in V and Z represent individual cells. Numbers in parenthesis in E, J, N indicate the number of subject animals. Numbers in parenthesis V and Z indicate the number of recorded cells. (J, N, V, Z) Paired t test. (E) One sample Wilcoxon test. All statistical tests are two-tailed. All p≤0.05 are specified. If not indicated, p> 0.05. See Supplementary Table 2 for detailed statistics.

**Figure S6.**
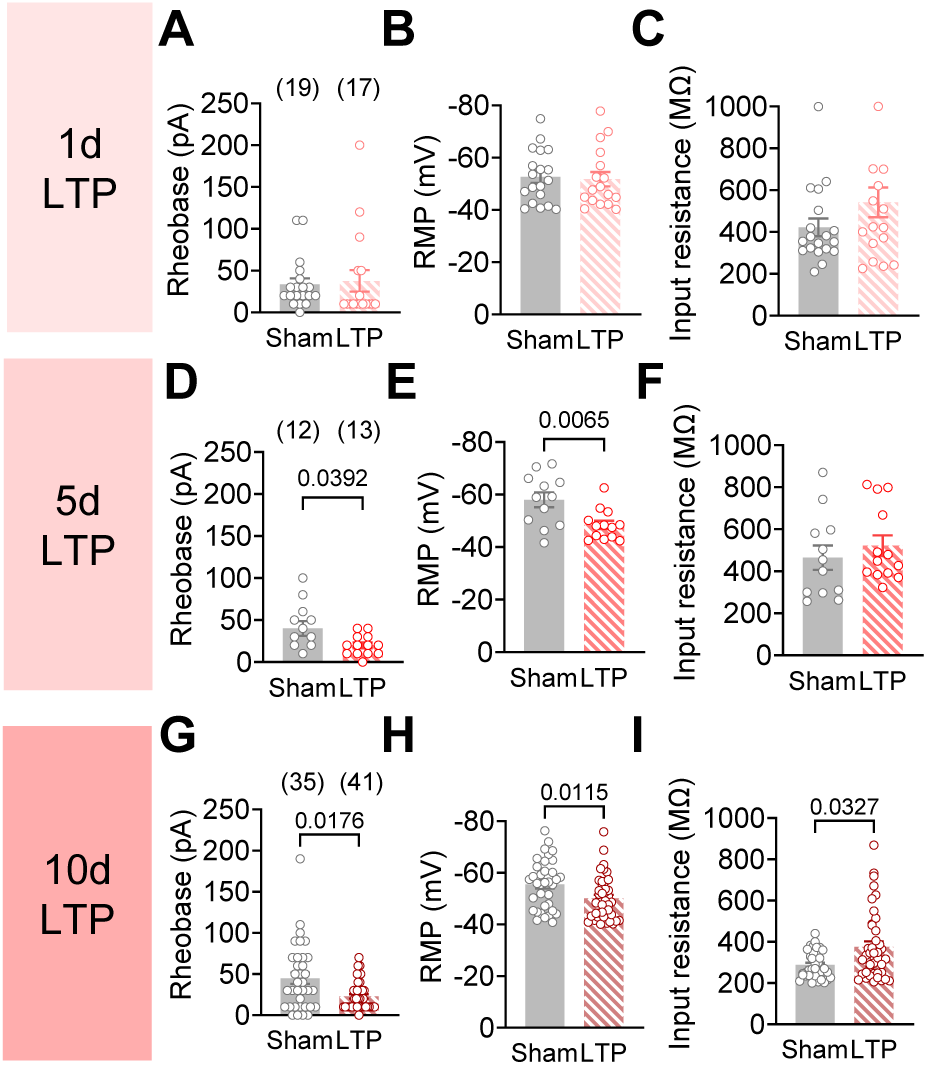
Intrinsic properties of cells on the sham and LTP side. Related to Figure 7. (A-C) The rheobase (A), resting potential (B), and input resistance (C) of VMHvl^Esr1^ cells on the sham and 1d LTP induction sides. (D-F) The rheobase (D), resting potential (E), and input resistance (F) of VMHvl^Esr1^ cells on the sham and 5d LTP induction sides. (G-I) The rheobase (D), resting potential (E), and input resistance (F) of VMHvl^Esr1^ cells on the sham and 10d LTP induction sides. Bars and error bars represent mean ± SEM. Circles represent individual cells. Numbers in parenthesis indicate the number of recorded cells. (B, D, E, H) Unpaired t test. (A, C, F, G, I) Mann-Whitney test. All statistical tests are two-tailed. All p≤0.05 are specified. If not indicated, p> 0.05. See Supplementary Table 2 for detailed statistics

**Figure S7.**
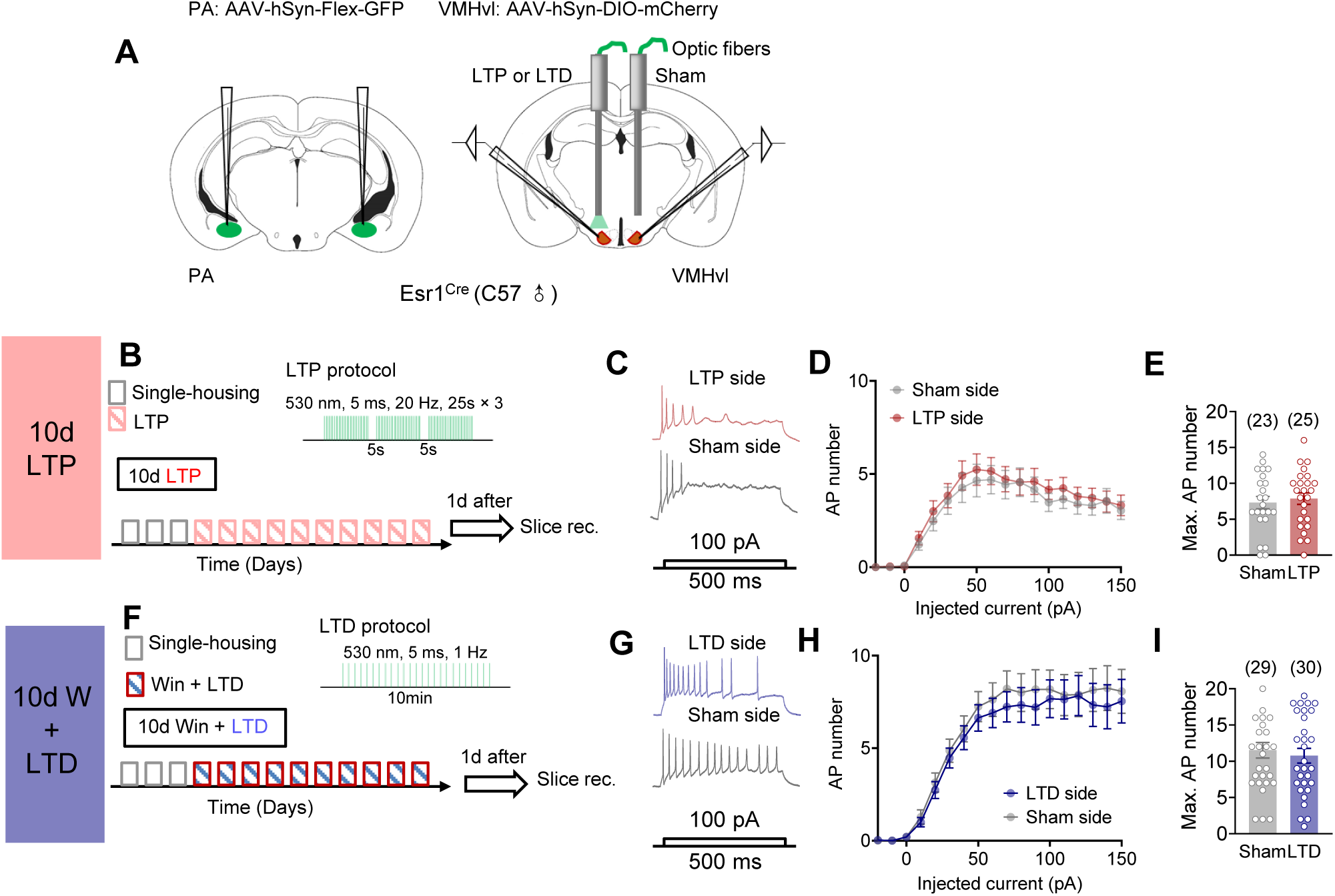
LTP and LTD induction protocol did not change VMHvl^Esr1^ cell excitability in control groups. Related to Figure 7. (A) Virus injection, fiber implantation, and slice recording strategies. (B) Experimental timeline. (C) Representative current clamp recording traces of VMHvl^Esr1^ cells on the sham and LTP light delivery sides with 100 pA current injection. (D) The F-I curve of cells recorded on the sham side and 10d LTP induction side. (E) Maximal number of action potentials across current steps of cells recorded from the sham and LTP induction sides. (F) Experimental timeline. (G) Representative current-clamp recording traces of VMHvl^Esr1^ cells on the sham and LTD induction sides with 100 pA current injection. (H) The F-I curve of cells recorded on the sham side and 10d LTD stimulation side. (I) Maximal number of action potentials across current steps of cells recorded from the sham and LTD induction sides. Bars and error bars represent mean ± SEM. Circles represent individual cells. Numbers in parenthesis indicate the number of recorded cells. (D, H) Two-way ANOVA with repeated measures followed by Sidak’s multiple comparisons test. (E, I) Unpaired t test. All statistical tests are two-tailed. All p ≤ 0.05 are specified. If not indicated p > 0.05. See Supplementary Table 2 for detailed statistics.

**Figure S8.**
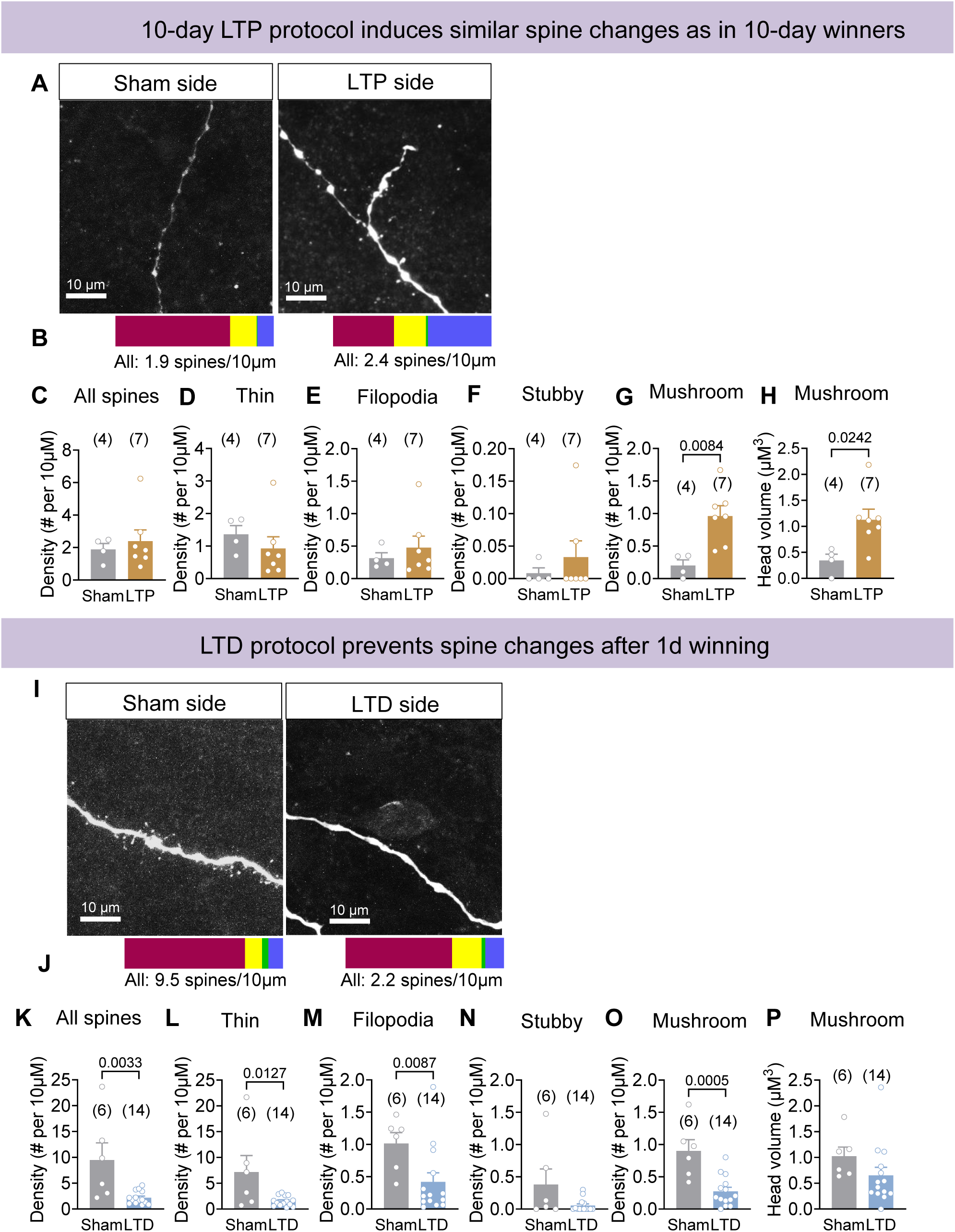
Spine morphology change in cells after LTP and LTD stimulation. Related to Figure 7. (A) Representative images of VMHvl^Esr1^ cell dendrites on the sham and 10d LTP stimulation sides. (B) The proportion of different types of spines on sham and 10d LTP sides. (C) Total spine density of cells on the sham and 10d LTP side. (D-G) The density of thin spines (C), filopodia (D), stubby spines (E), and mushroom spines (F) of cells on the sham and 10d LTP side. (H) The head volume of mushroom spines of cells on the sham and 10d LTP side. (I) Representative images of VMHvl^Esr1^ cell dendrites on the sham and LTD stimulation sides of a 1d winning mouse. (J) The proportion of different types of spines on sham and LTD sides. (K) Total spine density of cells on the sham and LTD sides of 1d winning mice. (L-O) The density of thin spines (L), filopodia (M), stubby spines (N), and mushroom spines (O) of cells on the sham and LTD sides of 1d winning mice. (P) The head volume of mushroom spines of cells on the sham and LTD sides. Bars and error bars represent mean ± SEM. Circles represent individual cells. Numbers in parenthesis indicate the number of cells. (B, C, D, F, I, J, M, N) Unpaired t test. (E, G, K, L) Mann-Whitney test. All statistical tests are two-tailed. All p≤0.05 are specified. If not indicated, p> 0.05. See Supplementary Table 2 for detailed statistics.

**Figure S9.**
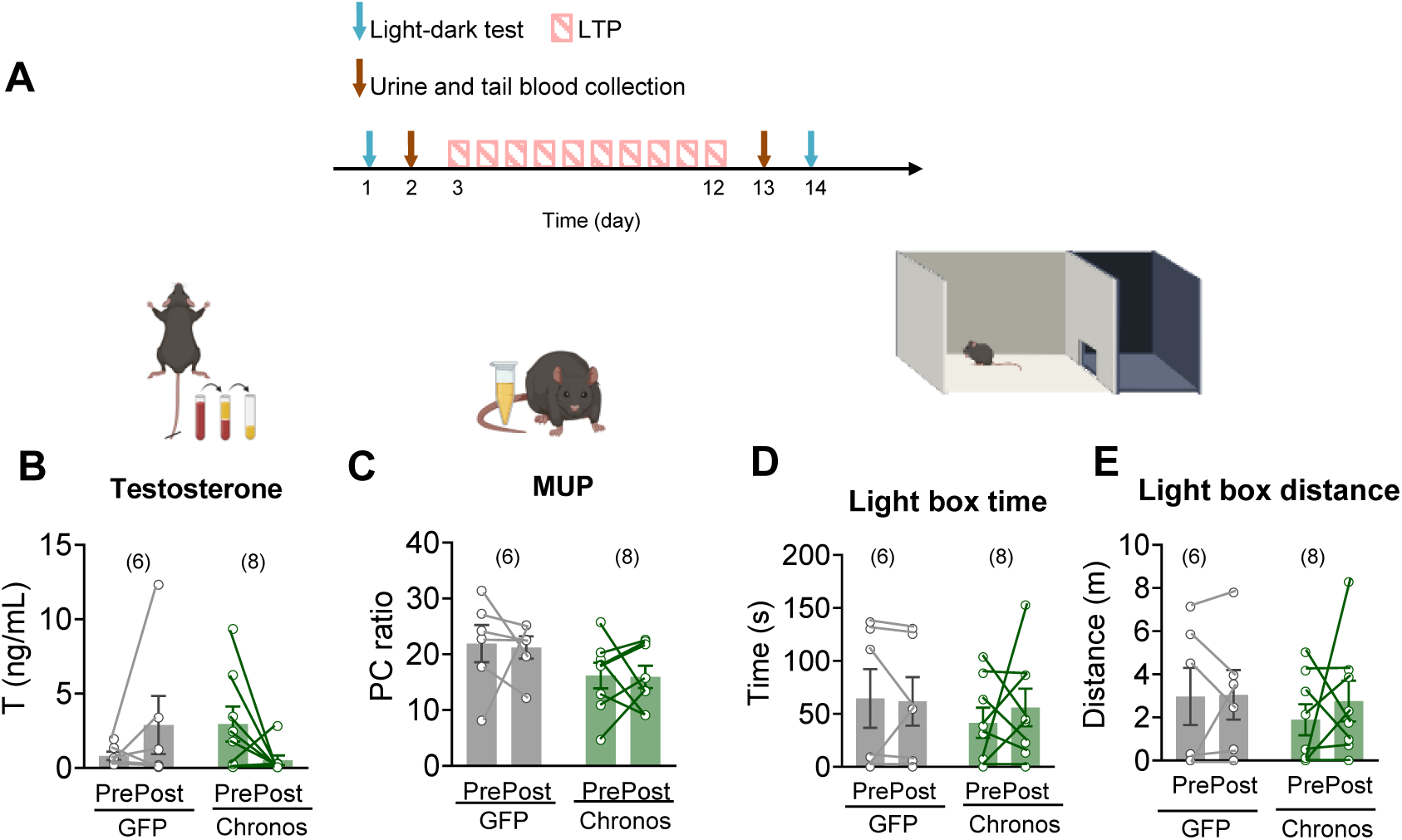
T and MUP levels and performance in the light-dark box do not change after 10d LTP induction. Related to Figure 7. (A) Timeline for urine/blood collection and the light-dark test in mice subjected to 10-day LTP induction. (B) The testosterone level before and after 10-day LTP stimulation in GFP- and Chronos-expressing mice. (C) The normalized MUP level (urine protein to creatinine ratio) before and after 10-day LTP stimulation in GFP- and Chronos-expressing mice. (D, E) The time spent (D) and distance traveled (E) in the light box before and after 10-day LTP stimulation. Bars and error bars represent mean ± SEM. Circles and lines represent individual animals. Numbers in parenthesis indicate the number of subject animals. (B, C, D, E) Two-way ANOVA with repeated measures followed by Sidak’s multiple comparisons test. All p ≤ 0.05 are specified. If not indicated, p > 0.05. See Supplementary Table 2 for detailed statistics.

## Notes

### Competing Interest Statement

The authors have declared no competing interest.

## References

Abdul-Ghani, M.A., Valiante, T.A., and Pennefather, P.S. (1996). Sr2+ and quantal events at excitatory synapses between mouse hippocampal neurons in culture. J Physiol 495 (Pt 1), 113–125.

Aizenman, C.D., and Linden, D.J. (2000). Rapid, synaptically driven increases in the intrinsic excitability of cerebellar deep nuclear neurons. Nature Neuroscience 3, 109–111.

Ammari, R., Monaca, F., Cao, M., Nassar, E., Wai, P., Del Grosso, N.A., Lee, M., Borak, N., Schneider-Luftman, D., and Kohl, J. (2023). Hormone-mediated neural remodeling orchestrates parenting onset during pregnancy. Science 382, 76–81.

Armano, S., Rossi, P., Taglietti, V., and D’Angelo, E. (2000). Long-term potentiation of intrinsic excitability at the mossy fiber-granule cell synapse of rat cerebellum. J Neurosci 20, 5208–5216.

Barkley, M.S., and Goldman, B.D. (1977). The effects of castration and Silastic implants of testosterone on intermale aggression in the mouse. Horm Behav 9, 32–48.

Batrinos, M.L. (2012). Testosterone and aggressive behavior in man. Int J Endocrinol Metab 10, 563–568.

Bekkers, J.M., and Clements, J.D. (1999). Quantal amplitude and quantal variance of strontium-induced asynchronous EPSCs in rat dentate granule neurons. J Physiol 516 (Pt 1), 227–248.

Calizo, L.H., and Flanagan-Cato, L.M. (2000). Estrogen selectively regulates spine density within the dendritic arbor of rat ventromedial hypothalamic neurons. J Neurosci 20, 1589–1596.

Chen, A.X., Yan, J.J., Zhang, W., Wang, L., Yu, Z.X., Ding, X.J., Wang, D.Y., Zhang, M., Zhang, Y.L., Song, N., et al. (2020). Specific Hypothalamic Neurons Required for Sensing Conspecific Male Cues Relevant to Inter-male Aggression. Neuron 108, 763–774 e766.

Chou, M.Y., Amo, R., Kinoshita, M., Cherng, B.W., Shimazaki, H., Agetsuma, M., Shiraki, T., Aoki, T., Takahoko, M., Yamazaki, M., et al. (2016). Social conflict resolution regulated by two dorsal habenular subregions in zebrafish. Science 352, 87–90.

Covington, H.E., 3rd, Newman, E.L., Leonard, M.Z., and Miczek, K.A. (2019). Translational models of adaptive and excessive fighting: an emerging role for neural circuits in pathological aggression. F1000Res 8.

Debanne, D., Inglebert, Y., and Russier, M. (2019). Plasticity of intrinsic neuronal excitability. Curr Opin Neurobiol 54, 73–82.

Demas, G.E., Munley, K.M., and Jasnow, A.M. (2023). A seasonal switch hypothesis for the neuroendocrine control of aggression. Trends Endocrinol Metab 34, 799–812.

Dias, I.C., Gutierrez-Castellanos, N., Ferreira, L., and Lima, S.Q. (2021). The Structural and Electrophysiological Properties of Progesterone Receptor-Expressing Neurons Vary along the Anterior-Posterior Axis of the Ventromedial Hypothalamus and Undergo Local Changes across the Reproductive Cycle. eNeuro 8.

Dreher, J.C., Dunne, S., Pazderska, A., Frodl, T., Nolan, J.J., and O’Doherty, J.P. (2016). Testosterone causes both prosocial and antisocial status-enhancing behaviors in human males. Proc Natl Acad Sci U S A 113, 11633–11638.

Falkner, A.L., Grosenick, L., Davidson, T.J., Deisseroth, K., and Lin, D. (2016). Hypothalamic control of male aggression-seeking behavior. Nat Neurosci 19, 596–604.

Frankfurt, M., Gould, E., Woolley, C.S., and McEwen, B.S. (1990). Gonadal steroids modify dendritic spine density in ventromedial hypothalamic neurons: a Golgi study in the adult rat. Neuroendocrinology 51, 530–535.

Franklin, T.B., Silva, B.A., Perova, Z., Marrone, L., Masferrer, M.E., Zhan, Y., Kaplan, A., Greetham, L., Verrechia, V., Halman, A., et al. (2017). Prefrontal cortical control of a brainstem social behavior circuit. Nat Neurosci 20, 260–270.

Ghani, M.U., Mesadi, F., Kanik, S.D., Argunsah, A.O., Hobbiss, A.F., Israely, I., Unay, D., Tasdizen, T., and Cetin, M. (2017). Dendritic spine classification using shape and appearance features based on two-photon microscopy. J Neurosci Methods 279, 13–21.

Grutzendler, J., Kasthuri, N., and Gan, W.B. (2002). Long-term dendritic spine stability in the adult cortex. Nature 420, 812–816.

Guo, Z., Yin, L., Diaz, V., Dai, B., Osakada, T., Lischinsky, J.E., Chien, J., Yamaguchi, T., Urtecho, A., Tong, X., et al. (2023). Neural dynamics in the limbic system during male social behaviors. Neuron 111, 3288–3306 e3284.

Hashikawa, K., Hashikawa, Y., Tremblay, R., Zhang, J., Feng, J.E., Sabol, A., Piper, W.T., Lee, H., Rudy, B., and Lin, D. (2017). Esr1(+) cells in the ventromedial hypothalamus control female aggression. Nat Neurosci 20, 1580–1590.

Holtmaat, A.J., Trachtenberg, J.T., Wilbrecht, L., Shepherd, G.M., Zhang, X., Knott, G.W., and Svoboda, K. (2005). Transient and persistent dendritic spines in the neocortex in vivo. Neuron 45, 279–291.

Hsu, Y., Earley, R.L., and Wolf, L.L. (2006). Modulation of aggressive behaviour by fighting experience: mechanisms and contest outcomes. Biol Rev Camb Philos Soc 81, 33–74.

Keppeler, D., Merino, R.M., Lopez de la Morena, D., Bali, B., Huet, A.T., Gehrt, A., Wrobel, C., Subramanian, S., Dombrowski, T., Wolf, F., et al. (2018). Ultrafast optogenetic stimulation of the auditory pathway by targeting-optimized Chronos. EMBO J 37.

Klapoetke, N.C., Murata, Y., Kim, S.S., Pulver, S.R., Birdsey-Benson, A., Cho, Y.K., Morimoto, T.K., Chuong, A.S., Carpenter, E.J., Tian, Z., et al. (2014). Independent optical excitation of distinct neural populations. Nat Methods 11, 338–346.

Kraeuter, A.K., Guest, P.C., and Sarnyai, Z. (2019). The Open Field Test for Measuring Locomotor Activity and Anxiety-Like Behavior. Methods Mol Biol 1916, 99–103.

Kudriavtseva, N.N., Bakshtanovskaia, I.V., and Avgustinovich, D.F. (1997). [The effect of the repeated experience of aggression in daily confrontations on the individual and social behavior of male mice]. Zh Vyssh Nerv Deiat Im I P Pavlova 47, 86–97.

Kudryavtseva, N.N. (2012). Psychopathology of Repeated (Animal) Aggression. In Encyclopedia of the Sciences of Learning, N.M. Seel, ed. (Boston, MA: Springer US), pp. 2731–2733.

Kudryavtseva, N.N., Bondar, N.P., and Alekseyenko, O.V. (2000). Behavioral correlates of learned aggression in male mice. 26, 386–400.

Kudryavtseva, N.N., Bondar, N.P., and Avgustinovich, D.F. (2004). Effects of repeated experience of aggression on the aggressive motivation and development of anxiety in male mice. Neurosci Behav Physiol 34, 721–730.

Kudryavtseva, N.N., Smagin, D.A., Kovalenko, I.L., and Vishnivetskaya, G.B. (2014). Repeated positive fighting experience in male inbred mice. Nat Protoc 9, 2705–2717.

Lee, H., Kim, D.W., Remedios, R., Anthony, T.E., Chang, A., Madisen, L., Zeng, H., and Anderson, D.J. (2014). Scalable control of mounting and attack by Esr1+ neurons in the ventromedial hypothalamus. Nature 509, 627–632.

Lee, J.-H., Kim, W.B., Park, E.H., and Cho, J.-H. (2023). Neocortical synaptic engrams for remote contextual memories. Nature Neuroscience 26, 259–273.

Lee, W., Khan, A., and Curley, J.P. (2017). Major urinary protein levels are associated with social status and context in mouse social hierarchies. Proc Biol Sci 284.

Lin, D., Boyle, M.P., Dollar, P., Lee, H., Lein, E., Perona, P., and Anderson, D.J. (2011). Functional identification of an aggression locus in the mouse hypothalamus. Nature 470, 221.

Lischinsky, J.E., and Lin, D. (2020). Neural mechanisms of aggression across species. Nature Neuroscience 23, 1317–1328.

Lo, L., Yao, S., Kim, D.W., Cetin, A., Harris, J., Zeng, H., Anderson, D.J., and Weissbourd, B. (2019). Connectional architecture of a mouse hypothalamic circuit node controlling social behavior. Proc Natl Acad Sci U S A 116, 7503–7512.

Lorenz, K. (1966). On aggression (London,: Methuen).

Matsuzaki, M., Ellis-Davies, G.C., Nemoto, T., Miyashita, Y., Iino, M., and Kasai, H. (2001). Dendritic spine geometry is critical for AMPA receptor expression in hippocampal CA1 pyramidal neurons. Nat Neurosci 4, 1086–1092.

Mei, L., Osakada, T., and Lin, D. (2023a). Hypothalamic control of innate social behaviors. Science 382, 399–404.

Mei, L., Yan, R., Yin, L., Sullivan, R.M., and Lin, D. (2023b). Antagonistic circuits mediating infanticide and maternal care in female mice. Nature 618, 1006–1016.

Nair, A., Karigo, T., Yang, B., Ganguli, S., Schnitzer, M.J., Linderman, S.W., Anderson, D.J., and Kennedy, A. (2023). An approximate line attractor in the hypothalamus encodes an aggressive state. Cell 186, 178–193 e115.

Ning, L., Suleiman, H.Y., and Miner, J.H. (2020). Synaptopodin Is Dispensable for Normal Podocyte Homeostasis but Is Protective in the Context of Acute Podocyte Injury. Journal of the American Society of Nephrology : JASN 31, 2815–2832.

Nishizuka, M., and Pfaff, D.W. (1989). Intrinsic synapses in the ventromedial nucleus of the hypothalamus: an ultrastructural study. The Journal of comparative neurology 286, 260–268.

Nordman, J.C., Ma, X., Gu, Q., Potegal, M., Li, H., Kravitz, A.V., and Li, Z. (2020). Potentiation of Divergent Medial Amygdala Pathways Drives Experience-Dependent Aggression Escalation. J Neurosci 40, 4858–4880.

Oyegbile, T.O., and Marler, C.A. (2005). Winning fights elevates testosterone levels in California mice and enhances future ability to win fights. Horm Behav 48, 259–267.

Parker, G.A. (1974). Assessment strategy and the evolution of fighting behaviour. J Theor Biol 47, 223–243.

Pchitskaya, E., and Bezprozvanny, I. (2020). Dendritic Spines Shape Analysis-Classification or Clusterization? Perspective. Frontiers in synaptic neuroscience 12, 31.

Petreanu, L., Mao, T., Sternson, S.M., and Svoboda, K. (2009). The subcellular organization of neocortical excitatory connections. Nature 457, 1142–1145.

Saito, K., He, Y., Yan, X., Yang, Y., Wang, C., Xu, P., Hinton, A.O., Jr., Shu, G., Yu, L., Tong, Q., et al. (2016). Visualizing estrogen receptor-α-expressing neurons using a new ERα-ZsGreen reporter mouse line. Metabolism: clinical and experimental 65, 522–532.

Shao, Y.Q., Fan, L., Wu, W.Y., Zhu, Y.J., and Xu, H.T. (2022). A developmental switch between electrical and neuropeptide communication in the ventromedial hypothalamus. Current biology : CB 32, 3137–3145.e3133.

Soden, M.E., Miller, S.M., Burgeno, L.M., Phillips, P.E.M., Hnasko, T.S., and Zweifel, L.S. (2016). Genetic Isolation of Hypothalamic Neurons that Regulate Context-Specific Male Social Behavior. Cell reports 16, 304–313.

Stagkourakis, S., Spigolon, G., Liu, G., and Anderson, D.J. (2020). Experience-dependent plasticity in an innate social behavior is mediated by hypothalamic LTP. Proc Natl Acad Sci U S A 117, 25789–25799.

Stagkourakis, S., Spigolon, G., Williams, P., Protzmann, J., Fisone, G., and Broberger, C. (2018). A neural network for intermale aggression to establish social hierarchy. Nat Neurosci 21, 834–842.

Takao, K., and Miyakawa, T. (2006). Light/dark transition test for mice. J Vis Exp, 104.

Tinbergen, N. (1951). The study of instinct (Oxford Eng.: Clarendon Press).

Wang, F., Zhu, J., Zhu, H., Zhang, Q., Lin, Z., and Hu, H. (2011). Bidirectional control of social hierarchy by synaptic efficacy in medial prefrontal cortex. Science 334, 693–697.

Wei, D., Osakada, T., Guo, Z., Yamaguchi, T., Varshneya, A., Yan, R., Jiang, Y., and Lin, D. (2023). A hypothalamic pathway that suppresses aggression toward superior opponents. Nature Neuroscience 26, 774–787.

Williamson, C.M., Lee, W., and Curley, J.P. (2016). Temporal dynamics of social hierarchy formation and maintenance in male mice. Animal Behaviour 115, 259–272.

Wong, L.C., Wang, L., D’Amour, J.A., Yumita, T., Chen, G., Yamaguchi, T., Chang, B.C., Bernstein, H., You, X., Feng, J.E., et al. (2016). Effective Modulation of Male Aggression through Lateral Septum to Medial Hypothalamus Projection. Current biology : CB 26, 593–604.

Yamaguchi, T., Wei, D., Song, S.C., Lim, B., Tritsch, N.X., and Lin, D. (2020). Posterior amygdala regulates sexual and aggressive behaviors in male mice. Nat Neurosci.

Yang, C.F., Chiang, M.C., Gray, D.C., Prabhakaran, M., Alvarado, M., Juntti, S.A., Unger, E.K., Wells, J.A., and Shah, N.M. (2013). Sexually dimorphic neurons in the ventromedial hypothalamus govern mating in both sexes and aggression in males. Cell 153, 896–909.

Yin, L., Hashikawa, K., Hashikawa, Y., Osakada, T., Lischinsky, J.E., Diaz, V., and Lin, D. (2022). VMHvllCckar cells dynamically control female sexual behaviors over the reproductive cycle. Neuron 110, 3000–3017.e3008.

Yoo, S., Cha, D., Kim, S., Jiang, L., Cooke, P., Adebesin, M., Wolfe, A., Riddle, R., Aja, S., and Blackshaw, S. (2020). Tanycyte ablation in the arcuate nucleus and median eminence increases obesity susceptibility by increasing body fat content in male mice. Glia 68, 1987–2000.

Yuste, R., and Bonhoeffer, T. (2004). Genesis of dendritic spines: insights from ultrastructural and imaging studies. Nat Rev Neurosci 5, 24–34.

Zhang, J., Dong, X.J., Ding, M.R., You, C.Y., Lin, X., Wang, Y., Wu, M.J., Xu, G.F., and Wang, G.D. (2020). Resveratrol decreases high glucose–induced apoptosis in renal tubular cells via suppressing endoplasmic reticulum stress. Molecular medicine reports 22, 4367–4375.

Zhang, W., and Linden, D.J. (2003). The other side of the engram: experience-driven changes in neuronal intrinsic excitability. Nature Reviews Neuroscience 4, 885–900.

Zhou, T., Zhu, H., Fan, Z., Wang, F., Chen, Y., Liang, H., Yang, Z., Zhang, L., Lin, L., Zhan, Y., et al. (2017). History of winning remodels thalamo-PFC circuit to reinforce social dominance. Science 357, 162–168.

Ziv, N.E., and Smith, S.J. (1996). Evidence for a role of dendritic filopodia in synaptogenesis and spine formation. Neuron 17, 91–102.

